# A machine learning method for calculating highly localized protein stabilities

**DOI:** 10.1101/2025.10.21.683809

**Authors:** Chenlin Lu, Kyle C. Weber, Savannah K. McBride, Andrew Reckers, Anum Glasgow

**Author notes:** Equal contributions (Chenlin Lu, Kyle C. Weber).

## Abstract

The residue-level free energy of opening (ΔG_op_) is the ultimate thermodynamic descriptor of localized protein stability, providing valuable information about the protein ensemble at physiologically relevant timescales and conditions. PFNet instantly determines ΔG_op_ for arbitrarily large proteins and complexes from conventional peptide-level hydrogen exchange/mass spectrometry (HX/MS) datasets. It unlocks the full potential of HX/MS, democratizing the method and establishing quantitative, scalable and accessible analysis (https://github.com/glasgowlab/PFNet).

## Main text

A complete understanding of protein function requires moving beyond static snapshots to the conformational ensembles that drive biological activity. Mutations, post-translational modifications, and molecular interactions regulate protein functions by reweighting their ensembles. Thus, probing shifts in the conformational ensembles of proteins across physiologically relevant timescales and conditions is essential for truly understanding the molecular basis for protein function and enabling the rational design of sophisticated functional proteins. However, a fundamental technological barrier has hindered this pursuit: the lack of tools to quantitatively monitor these energetic changes at high resolution.

Hydrogen exchange/mass spectrometry (HX/MS) is a powerful technique that monitors the ensemble-averaged kinetics of hydrogen isotope exchange in the protein backbone, thus reporting on localized protein stabilities. The data are typically interpreted qualitatively via plots that show relative differences in peptide deuterium uptake among protein functional states. However, in theory, HX/MS data contain all the information necessary to derive individual hydrogen exchange rates (k_ex_) for most backbone amide groups in a protein, which can be used to calculate residue-level free energies of opening (ΔG_op_)^1–3^: the ultimate thermodynamic descriptor of the localized stability.^4^ Despite recent advances^3–5^, a method that simultaneously achieves accuracy, efficiency, scalability, and user-friendliness remains elusive, thereby impeding the full potential of ΔG_op_ as a scalable tool for protein ensemble analysis.

We therefore developed PFNet, a machine learning model with a transformer architecture that can determine single-residue ΔG_op_ from conventional peptide-level HX/MS datasets within seconds. PFNet takes isotopic mass envelopes from HX/MS experiments on proteins as inputs, encodes them as timepoint token representations, and outputs residue-level log(k_ex_) at the highest spatial resolution that the data afford, with confidence scores (Fig. 1a). We trained PFNet on more than two million proteins with 1,000-10,000 synthetic HX/MS isotopic envelopes each. The training data mimicked real HX/MS data by including varied timepoints, noise, peptide coverage, and D-to-H back-exchange, which occurs in the chromatography steps of the experiment. Because most published HX/MS datasets only include MS centroids rather than full isotopic mass envelopes, we also developed the alternative model PFNet-Centroid.

**Figure 1.**
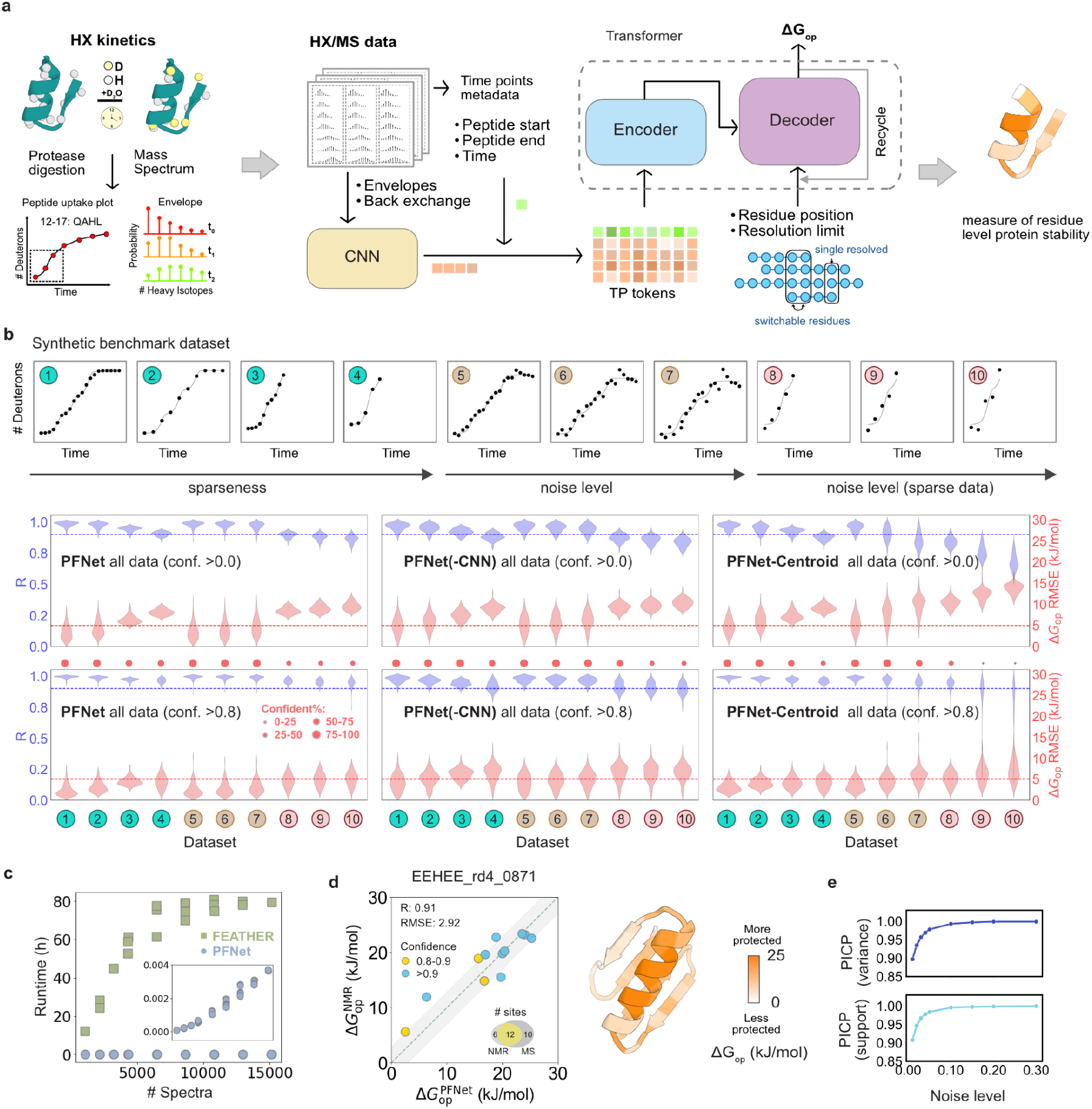
PFNet architecture and benchmarking. **a**, Schematic representation of the PFNet architecture and data flow. **b**, Benchmarking of PFNet, PFNet(–CNN), and PFNet-Centroid on synthetic datasets with varying time windows, sparsity, and noise levels (Table S1), illustrated by a schematic representation of a peptide uptake plot. The Pearson correlation coefficient (R) and root-mean-squared error (RMSE) distributions were calculated with and without applying a confidence threshold of 0.8, respectively. **c**, Runtime comparison between PFNet and FEATHER on the *perfect* synthetic dataset (Table S1). **d**, Experimental benchmarking on the EEHEE_rd4_0871 dataset. The Venn diagram shows the number of single-residue sites resolved by each technique. **e**, PICP analysis estimating the equivalent noise levels of the EEHEE_rd4_0871 HX/MS dataset corresponding to the synthetic benchmarks.

We first developed a synthetic HX/MS benchmark dataset to test PFNet performance and runtime without the confounding factors present in experimental datasets (Fig. 1b). The benchmark set comprises 1000 synthetic proteins that closely resemble experimental observations in terms of quality, noise, redundancy, and spatiotemporal sparseness (Table S1). We used two metrics to assess model performance: the Pearson correlation of the log(k_ex_) predictions to the true log(k_ex_) (R); and the root-mean-squared error (RMSE).

Testing PFNet on the benchmark data revealed that it achieves higher accuracy and greater robustness to noise than FEATHER (Fig. 1b, Supplemental Figs. S1, S2), a state-of-the-art Bayesian method for high-resolution ΔG_op_ determination^3^, but with four to five orders of magnitude less analysis time (Fig. 1c, from tens of hours to seconds). Further, PFNet benefits considerably from increased numbers of peptides, demonstrating strong data efficiency and scalable performance (Supplemental Fig. S1). By contrast, PFNet-Centroid suffers from reduced accuracy due to information loss from centroiding the data (Fig. 1b, Supplemental Fig. S3).^6^ Ablating the convolutional neural network (CNN) that pre-encodes isotopic mass envelopes markedly reduced predictive accuracy, highlighting the essential roles of both envelope information and CNN-based encoding (Fig. 1b, Supplemental Fig. S4). The final PFNet model embeds this CNN-encoded envelope information within its architecture to achieve optimal performance. When predictions were filtered based on their confidence scores, the RMSE was reduced, indicating that the confidence score is a reliable proxy for prediction accuracy, particularly in sparse and noisy datasets. The full PFNet model achieves the highest proportion of confident predictions (residues with confidence >0.8, see red dot size in Fig. 1b), providing both the most confident and most accurate results. Although PFNet-Centroid shows comparatively reduced overall performance, its confident predictions remain reliable.

Satisfied with the theoretical accuracy of PFNet, we then compared its predictions to the experimental HX/NMR-derived ΔG_op_ for two proteins: a small *de novo-*designed mini-protein (Fig. 1d, Supplemental Figs. S5-S7, Tables S2, S3), and T4 lysozyme (T4L) (Supplemental Figs. S8-10, Tables S4, S5).^7,8^ Although HX/NMR can determine residue-level ΔG_op_, only about 100 datasets are available due to technical limitations. HX/NMR is ideal for small proteins in a narrow pH range that are isotopically labeled and purified, and it can take months to make the resonance assignments. For both the mini-protein and T4L, PFNet produced a set of ΔG_op_ that closely matched those determined by HX/NMR (Fig. 1d, Supplemental Fig. S9). To estimate the noise level inherent in the experimental HX/MS datasets in comparison with the benchmarks, we used the prediction interval coverage probability (PICP)^9^ relative to the theoretical intervals of the synthetic noise added during training (Fig. 1e, see Methods). The variance equivalent interval measures how well the simulated noise reproduces the variance of the experimental residuals, whereas the support interval compares the maximum range of the possible deviations. The equivalent noise level was defined as the point at which PICP reached the nominal 0.95 coverage. The estimated noise levels of the mini-protein (Fig. 1e) and T4L (Supplemental Fig. S11) are 0.03 and 0.04 respectively, suggesting that the variation of these datasets is low. This correspondence allows the expected prediction accuracy for datasets to be inferred directly from the synthetic benchmarks (Fig. 1b, Supplemental Fig. S11), in agreement with the NMR benchmarks. Additionally, PFNet is robust not only to noise in the synthetic benchmarks but also in experimental datasets, with additional noise injections showing that prediction accuracy remained stable (Supplemental Fig. S7).

HX/MS can reveal how a protein’s ensemble responds to molecular perturbations like ligand binding and mutations. To determine how well PFNet captured energetic differences between protein functional states (ΔΔG_op_), we compared its performance to FEATHER for apo and trimethoprim (TMP)-bound *E. coli* dihydrofolate reductase (ecDHFR). TMP is an antibiotic that stabilizes a distinct conformational state of ecDHFR.^3^ PFNet recapitulated the FEATHER results (Fig. 2a, Supplemental Fig. S12) with lower fitting errors (Fig. 2b, Supplemental Fig. S13), which suggests that the PFNet-derived ΔG_op_ values more closely match the experimental observations. Notably, the predictions of PFNet-Centroid showed good correlation with those of PFNet, as demonstrated by the synthetic dataset results (Fig. 2c).

**Figure 2.**
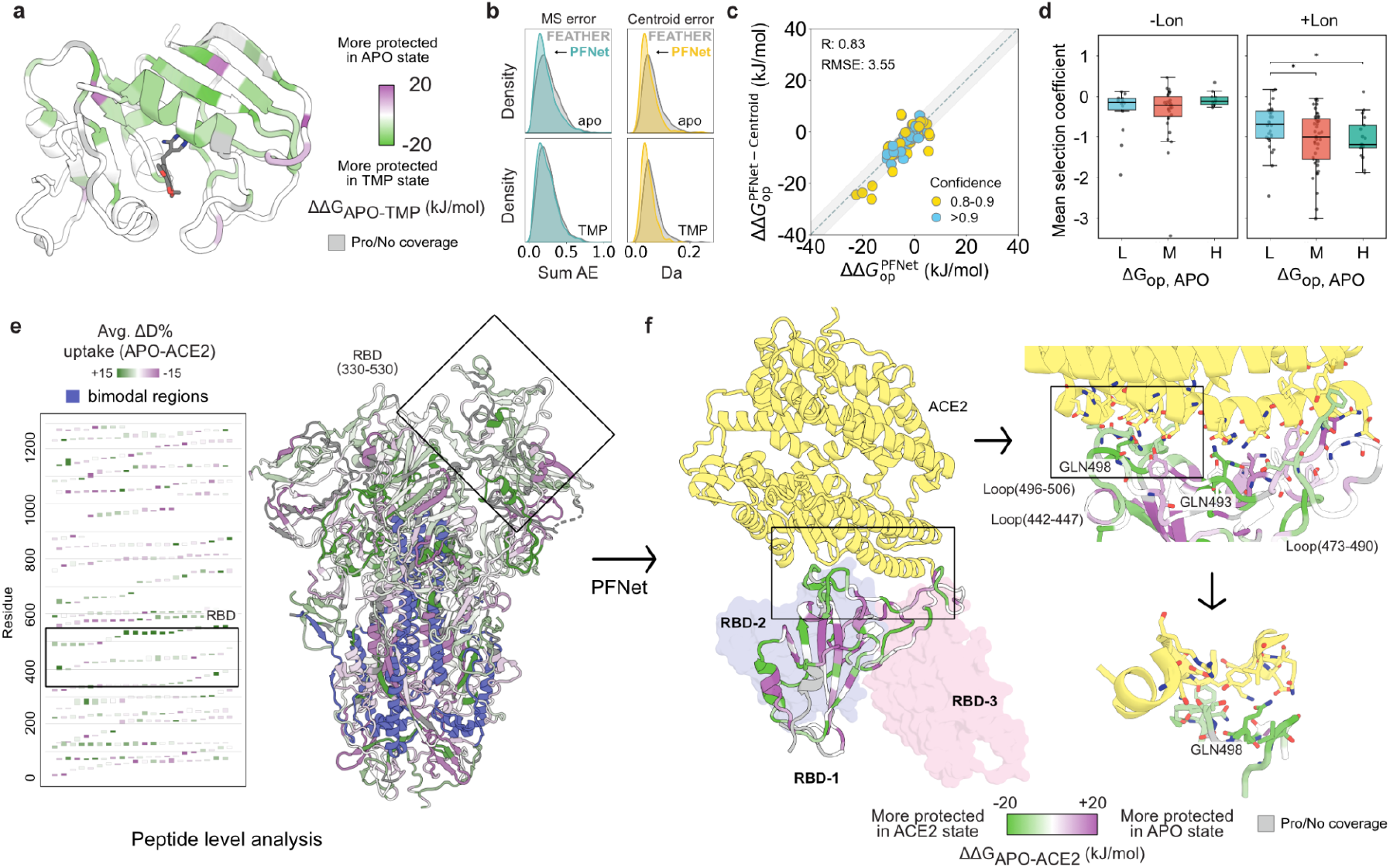
PFNet reveals how therapeutic targets respond to binding partners. **a**, PFNet-derived ΔΔG_op(apo-TMP)_ values mapped onto the crystal structure of ecDHFR (PDB: 6XG5). **b**, Comparison of fitting errors between PFNet and FEATHER-derived results on the ecDHFR dataset, shown as distributions of summed absolute errors for mass spectra and absolute fitting errors for centroids). **c**. Correlation between ΔΔG_op(apo-TMP)_ values predicted by PFNet and PFNet-Centroid for ecDHFR. **d**. Distribution of selection coefficients across three ΔG_op(apo)_ groups in the absence or presence of Lon protease from DMS (Thompson *et al*.), sorted by ΔG_op(apo)_ as defined by the 25th and 75th percentiles: Low (< 9.5 kJ mol−^1^), Medium (9.5– 24.1 kJ mol−^1^), and High (> 24.1 kJ mol−^1^). Asterisks indicate significant pairwise differences determined by the Mann–Whitney U test (p < 0.05). **e**, Peptide-level analysis of deuterium uptake differences in the SARS-CoV-2 spike protein between the apo and ACE2-bound states. Bimodal regions are shown in blue (Supplemental Fig. S15). **f**, PFNet-derived ΔΔG_op_ profiles enable high-resolution, unbiased thermodynamic epitope mapping.

Interestingly, in comparing the apo ecDHFR PFNet results with a deep mutational scanning (DMS) dataset collected by Thompson *et al*.^10^, we find that highly protected residues show significant drops in DMS selection coefficients upon mutation in the presence of the global regulatory protease Lon but remain comparatively tolerant to mutation in its absence (Fig. 2d). This indicates that mutations in the most locally stable regions of ecDHFR become significantly less tolerated when Lon protease is active, reflecting increased sensitivity to local destabilization and partial unfolding. Thus, understanding residue-level ΔG_op_ provides a thermodynamic basis for predicting mutational tolerance, offering valuable guidance for protein stability engineering and directed evolution.

To investigate how PFNet might illuminate viral protein-host interactions, we applied the method to the SARS-CoV-2 spike protein ectodomain interaction with the human ACE2 receptor, using an HX/MS dataset collected on a different mass spectrometer by another group (Tables S6, S7).^11^ The spike-ACE2 interaction is the first step of viral cell invasion. Notably, such large complexes are intractable to HX/NMR. Comparing the peptide-level qualitative analysis with the residue-level PFNet analysis (Fig. 2e, f, Supplemental Fig. S14), we found that the PFNet-derived ΔΔG_op_ values enable high-resolution and unbiased thermodynamic epitope mapping, providing quantitative insights that are inaccessible to conventional peptide-level HX/MS. This analysis revealed that loops 497–506 and 442–447 are more strongly protected than loop 473–482, whereas loop 473–482 shows partial destabilization. Among all residues, Gln498 and Gln493 exhibit the strongest protection, pinpointing these as the thermodynamic centers of ACE2 recognition (Fig. 2f). These findings are consistent with prior structural^12,13^ and mutational studies^14,15^: X-ray crystal structures show that Gln498 and Gln493 form key hydrogen bonds with ACE2^12,13^, which informed the design of ACE2 receptor traps as therapeutics^16^; DMS demonstrated that substitutions of Gln498 and Gln493 alter ACE2 affinity, defining them as a functional hotspots^14,15^; and molecular dynamics simulations revealed a long-lived contact between Gln498 and ACE2 Lys353^17^. In the Omicron variant of SARS-Cov2, RBD mutations Q493R and G496S further increase the binding affinity via new hydrogen bonds with ACE2 Glu35 and Lys353.^18^ Together, these observations reinforce the roles of Gln498 and Gln493 as dominant contributors to ACE2 binding stability.

Moreover, the ΔΔG_op_ profile derived from PFNet revealed coupled thermodynamic changes upon ACE2 binding. The ACE2 binding-induced population shift toward the RBD “up” conformation^19,20^ leads to increased protection and rigidity in several loop regions (Supplemental Fig. S14). This local stabilization is accompanied by destabilization of the central β-sheet, suggesting an enthalpy-entropy compensation mechanism^21,22^ that preserves the overall spontaneity of ACE2 binding (Fig. 2f). Structural comparisons between the RBD “up” and “down” conformations support this interpretation, showing that apart from increased electron density in loop regions, the global fold remains nearly identical (root-mean-square deviation of 1.42 Å).^23,24^ The quantitative HX/MS pipeline also enabled the identification of additional bimodal regions to those previously reported^11^; these regions have been shown to report on two slowly-interconverting conformations of the same secondary structures that exist simultaneously in solution (Supplemental Fig. S15). These results show that PFNet resolves the precise energetic coupling between local protection and global flexibility that cannot be captured by static structural analyses alone, even in large, dynamic complexes.

HX/MS experiments should be designed towards optimizing the data analysis. An ideal dataset for PFNet is easy to collect: three pooled technical replicates, each with different timepoints and proteases, alongside fully deuterated controls for back-exchange correction and tandem MS experiments for peptide identification. The inclusion of data points across a large time window spanning seconds to 24+ hours is more valuable than dense data points in a narrow time window. Data collection and analysis can be completed in 1-2 days, with less than an hour of hands-on instrument time if the experiment is automated.^25^

While HX/MS remains popular for qualitative epitope-mapping applications, here we report how coupling PFNet with a standard HX/MS workflow can reveal detailed biomolecular mechanisms, making full use of the information-rich data without complicating the experiment. The method is transformative in protein science and engineering: compared to FEATHER and other methods for high-resolution ΔG_op_ determination, PFNet is more accurate with less data, dramatically faster, and easier to use, thereby democratizing HX/MS data collection and analysis such that all labs can use the method to extract the maximum amount of information about protein ensembles. The increased accuracy afforded by PFNet allows for the detection of subtle but crucial energetic differences among protein functional states within a single exceptionally sensitive framework, enabling a detailed understanding of biological machines.

PFNet can be applied to hundreds or thousands of proteins simultaneously, or to proteins with hundreds or thousands of residues, to understand their free energy landscapes and to generate large and accurate datasets for future machine learning applications. By providing a high-resolution energetic blueprint for protein interactions or mutational effects, PFNet may enable the precise rational design of allosteric therapeutics. The SI describes how to access the benchmark and experimental HX/MS datasets, convert HX/MS data for compatibility with PFNet, and use PFNet locally or on the cloud-based public installation.

## Methods

### Model

PFNet was built using the deep-learning Python packages PyTorch^26^ and PyTorch-Lightning^27^. PFNet employs a conventional encoder–decoder transformer architecture tailored for peptide-level HX/MS data. For each timepoint of each peptide, the isotopic mass envelope is first processed by a one-dimensional convolutional neural network (1D-CNN), which captures local spectral features. The resulting embeddings are concatenated with time encodings and peptide positional encodings to form a unified timepoint token representation. These timepoint tokens are subsequently passed through the transformer encoder, which models the temporal and spatial dependencies across all peptides and timepoints. The decoder queries consist of residue-level representations, incorporating information about intrinsic exchange rates (k_int_, the exchange rate for an unfolded peptide) and the resolution limits imposed by the peptide coverage of the dataset. PFNet outputs both the predicted residue-level log(k_ex_) values and their associated confidence scores by separate output heads using the same decoder embeddings. The confidence head is trained by mapping the predicted confidence values to the predicted absolute error using a single exponential decay function. The decoder also employs an iterative recycling mechanism, in which predicted outputs from the previous iteration are incorporated into the query embeddings, and the corresponding reconstruction errors are included as part of the key embeddings. This design allows the model to iteratively refine its predictions by leveraging prior outputs and their associated reconstruction errors. These predicted log(k_ex_) are subsequently used to compute the ΔG_op_.

### Loss function

We designed a composite loss function that balances predictive accuracy and physical consistency during model training:

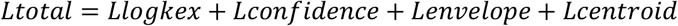

Here, *L_log_kex* is the mean squared error (MSE) between predicted and true residue-level log(k_ex_); *L_confidence* encourages well-calibrated uncertainty estimates; and *L_envelope* and *L_centroid* compare the reconstructed and input peptide-level isotopic envelopes and centroids. Envelope reconstruction is performed using the same forward calculation as previously reported given the predicted exchange rates.^3^ Loss weights are adjusted dynamically during training to emphasize different objectives at different stages.

### Synthetic training datasets

Synthetic datasets were generated using an established method.^3^ These datasets span a wide range of conditions, including protein sizes from 100 to 400 residues, peptide number ranging from 0.7 to 1.3 times the protein length, peptide lengths between 3 and 48 residues, labeling time windows from 10−^6^ to 10^18^ seconds and back exchange from 0.0 to 0.8, so as to mimic the range of conditions observed in experimental datasets, such as the ecDHFR data previously reported by our laboratory.^3^ To better approximate realistic experimental variability, diverse noise sources were incorporated, including additive noise to isotopic envelopes, back exchange, and side-chain exchange effects, all scaled by a global parameter (noise level). Back exchange levels were perturbed with Gaussian noise proportional to noise level/5. At non-zero time points, side-chain exchange noise was added as a small positive offset up to noise level/10 of the theoretical maximum uptake. Isotopic envelopes were further perturbed by baseline fluctuations proportional to both noise level and peak intensity, together with a time-dependent term scaled by noise level/10 × log10(t). All envelopes were truncated to non-negative values and renormalized. The noise level ranges from 0.1 to 0.3 in the training. Synthetic data were generated continuously in an online manner during training to ensure diversity and improve model generalization.

### Synthetic benchmarking datasets

Benchmarking datasets were generated in the same manner as the synthetic training datasets, with controlled protein size, peptide number, time windows, and noise levels to assess their effects on model performance (Table S1). For each dataset, five different proteins were generated to ensure adequate diversity and statistical robustness. Note: the noise sources differ from those used in the benchmarking dataset for PIGEON-FEATHER.^3^ We therefore add the prefix PFsim to indicate this distinction.

### Training

PFNet training was conducted in two stages. The *Llogkex* was applied throughout both phases. In phase 1, only the MSE and reconstruction losses (envelope and centroid) were used, while the confidence loss was turned off. This allowed the model to focus on learning accurate log(k_ex_) and reproducing peptide-level envelopes with respect to the HX physics. In phase 2, the confidence loss was enabled and the weights of reconstruction losses were reduced to avoid overfitting on the artificial noise, shifting the focus to predictive accuracy and uncertainty calibration. Training was performed on two NVIDIA RTX 4090 GPUs for two weeks. The training dataset comprised over 100 million synthetic protein examples, corresponding to approximately 10 billion simulated mass spectra.

### Back exchange modeling

There are two common approaches for modeling back exchange in the forward HX calculations, which convert residue level log(k_ex_) values into peptide-level isotopic mass envelopes. The first approach assigns a uniform back exchange value to each residue, independent of its peptide sequence context.^28^ The second approach distributes the peptide-level measured back exchange evenly across residues within a peptide.^3^ Both methods rely on a widely used assumption that the first or first and second residues of a peptide do not retain deuterium, *i*.*e*., their back exchange is 100%. However, this assumption is not universally valid for the second residue, which heavily depends on the sequence context.^29^ HX/NMR experiments show that back exchange behavior is primarily governed by k_int_, the intrinsic exchange rates of residues in the unfolded state, which are influenced by both solution conditions and the peptide sequence. For example, when a peptide begins with bulky residues such as “PI,” the second residue “I” significantly contributes to deuterium retention. In contrast, internal sequence motifs like “HH” can exhibit complete back exchange at the second “H” residue.^29,30^ These observations highlight that residue-specific back exchange is heterogeneous and sequence-dependent.

To incorporate such context dependent heterogeneity into back exchange modeling, we introduce an equivalent back exchange time under ideal quench conditions (pH 2.3, 0 °C). This time parameter is fitted to reproduce the experimentally measured back exchange for a given peptide and is then used to infer residue level back exchange values within that peptide. This approach provides a physically motivated, sequence aware refinement to conventional back exchange corrections.

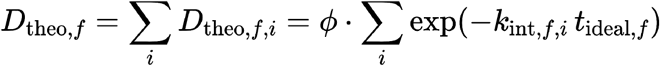

where D_{theo,f,i} denotes the theoretical deuterium retention of residue *i* in peptide *f* at t_ideal,f. k_{int,f,i} is the intrinsic exchange rate of residue *i* in peptide *f* under ideal quench conditions (pH 2.3, 0 °C), calculated using HDXrate, saturation level *ϕ* is the deuterium percent of the D_2_O buffer. The parameter t_{ideal,f} is determined using the brentq root-finding algorithm (scipy.optimize.brentq) to match the experimentally measured fully deuterated value for peptide *f*. Then the back exchange value of residue *i* in the context of peptide *f* β_f,i_ is

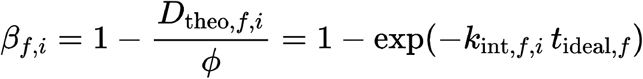

### Estimation of equivalent experimental noise level

To estimate the experimental noise level that is consistent with synthetic data, residuals were compared against the theoretical intervals of the synthetic two uniform noise used during training. We computed the prediction interval coverage probability (PICP) using two complementary interval definitions:

1. Variance equivalent interval, which reflects agreement in terms of variance.

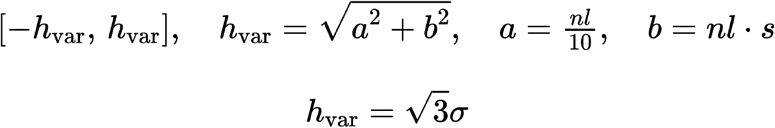 Here, *a* and *b* are the two uniform noise added during the training scale by the noise level.
2. Support interval, which provides a complementary check based on the maximum possible error range.

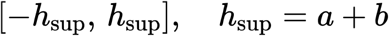

The noise level at which the PICP reached the nominal 0.95 coverage was defined as the equivalent experimental noise level, thereby aligning experimental variation with synthetic benchmarking conditions.

### HX/MS data collection and analysis

For T4L, we introduced a T4 bacteriophage cysteine-free gene with an N-terminal 6X-histidine tag into a modified pET9a plasmid via Golden Gate cloning, using a BsaI restriction endonuclease to produce the pET9a-6xHis-T4L plasmid. We chemically transformed the pET9a-6xHis-T4L plasmid into a BL21 LOBSTR strain. Cells were grown at 37 °C overnight shaking at 200 rpm in LB media with 50 µg/ml kanamycin, then subcultured 1:100 into 0.5 L LB media with 50 µg/ml kanamycin and grown at 30 °C until the optical density at 600 nm was measured between 0.2 and 0.4. Protein expression was induced by the addition of 1 mM IPTG and the cells were grown at 16 °C for 12-16 hours. The culture was centrifuged at 6,000×*g* for 20 minutes, and the cell pellet was frozen at -20 °C. After several weeks, the pellet was thawed in 20 ml lysis buffer: 50 mM Tris, 150 mM NaCl, 0.5 mM TCEP, pH 8.0 with 5 mM MgCl2, 1 mM MnCl2, 100 µM CaCl2, a protease inhibitor cocktail (Pierce), and 2000 units/ml DNaseI. The mixture was vortexed at 20 °C for 15 minutes, then sonicated in an ice bath at 30% amplitude for 10 minutes, with a four second break following every two seconds of sonication to prevent overheating. Next, the lysate was centrifuged at 27,000×*g* for 30 minutes at 4 °C. The soluble fraction was applied to 2 ml Ni-NTA resin (Thermo Fisher) in 5 ml sample buffer (50 mM Tris, 150 mM NaCl, 0.5 mM TCEP, pH 8.0), and the mixture left to nutate overnight at 4 °C. The mixture was then added to a 20 ml benchtop gravity column, allowed to drain and settle, and washed with 50 ml wash buffer (50 mM Tris, 250 mM NaCl, 0.5 mM TCEP, 20 mM imidazole, pH 8.0). The protein was eluted in 14 ml elution buffer (50 mM Tris, 150 mM NaCl, 0.5 mM TCEP, 200 mM imidazole, pH 8.0) and then purified further on a HiLoad16/600 Superdex 200 size exclusion column (Cytiva). The eluent at the retention time corresponding to the monomeric molecular weight was concentrated to 0.5 mg/ml using a 15 ml ultracentrifugal filter column with a 10 KDa molecular weight cutoff (MilliporeSigma). This purified stock was split into aliquots, flash frozen in liquid N2, and stored at -80 °C. On the day of HX/MS data collection and analysis, the protein stocks were thawed, and the 25 µM sample was diluted tenfold into the exchange buffer (50 mM Tris, 150 mM NaCl, 0.5 mM TCEP, pH 8.0 in D_2_O). Details for the data collection and analysis for T4L, ecDHFR, and the mini-protein, as well as the expression and purification of ecDHFR, were as previously described.^3^

### Merging T4 lysozyme datasets collected at two pHs

To obtain a broader exchange time window, we combined seven replicate datasets collected at pH 8.0 and pH 6.0 (Table S2). The measured timepoints at pH 8.0 were converted to their equivalents at pH 6.0 using the method described by Hamuro *et al*.^31,32^, which has been shown to effectively extend the time window range of exchange measurements for proteins that exhibit similar behavior across these pH conditions (Supplemental Fig. S8).

## Data availability

The data are provided in HXMS format^33^, which includes the isotopic mass envelope peaks or centroids for all timepoints and peptides as defined by starting and ending residues, as well as important experimental conditions that influence k_ex_ determination by the Linderstrøm-Lang model^34^: pH, D_2_O saturation level, and temperature. PFLink [https://huggingface.co/spaces/glasgow-lab/PFLink] enables the conversion of HX/MS data from commonly used software packages.^33^ The synthetic benchmark dataset is available at: [https://zenodo.org/records/17353731]. The raw HX/MS data for T4 lysozyme is available at the PRIDE database (identifier PXD069693).

## Code availability

Code for dataset generation, model training, inference, and access to trained models is available at: https://github.com/glasgowlab/PFNet. To facilitate broader accessibility, the trained model is also hosted on HuggingFace at https://huggingface.co/spaces/glasgow-lab/PFNet.

## Acknowledgements

We thank members of the Glasgow Lab for requesting, discussing, and testing various features of the model and analysis code. We are grateful to Dr. Gabriel Rocklin and his group for providing the *EEHEE_rd4_087* sample and NMR dataset, as well as for insightful discussions regarding these data, and to Drs. Susan Marqusee, Shawn Costello, and Sophie Shoemaker for sharing their SARS-CoV-2 spike/ACE2 raw data files. We also thank Dr. Mohammed AlQuraishi at Columbia University Medical Center for valuable discussions.

## Notes

https://github.com/glasgowlab/PFNet

https://huggingface.co/spaces/glasgow-lab/PFNet

### Competing Interest Statement

The authors have declared no competing interest.

### Funding

- National Institutes of Health: R35GM157185, R21EB035208

