## Supplemental information for "A machine learning method for calculating highly localized protein stabilities"

### 1. Supplemental figures

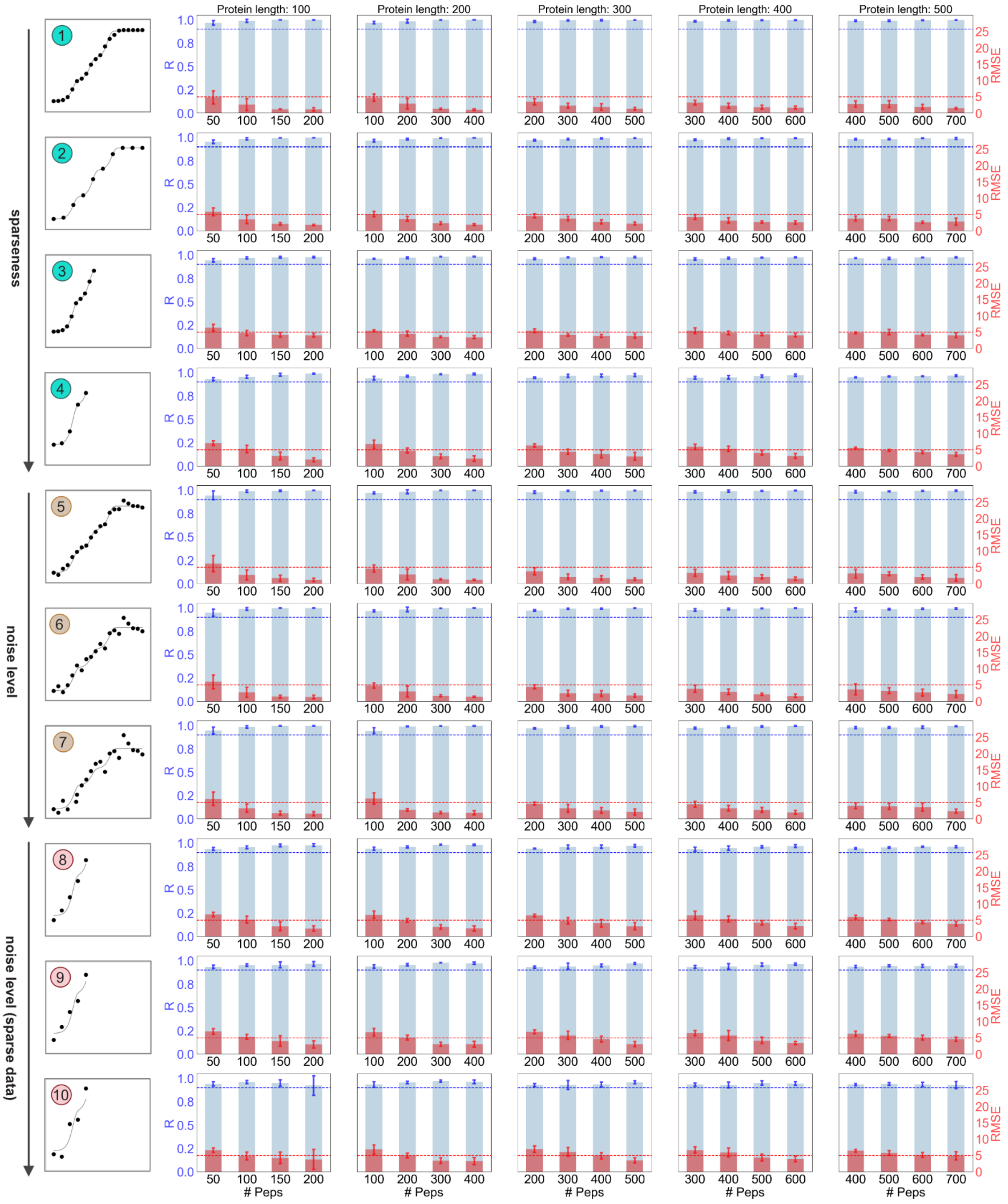

**Figure S1. Expanded data of Fig. 1b for PFNet predictions with confidence > 0.8.**

PFNet performance on synthetic datasets with varying degrees of sparsity and noise across proteins of different sizes and peptide numbers, with R (blue) and RMSE (red).

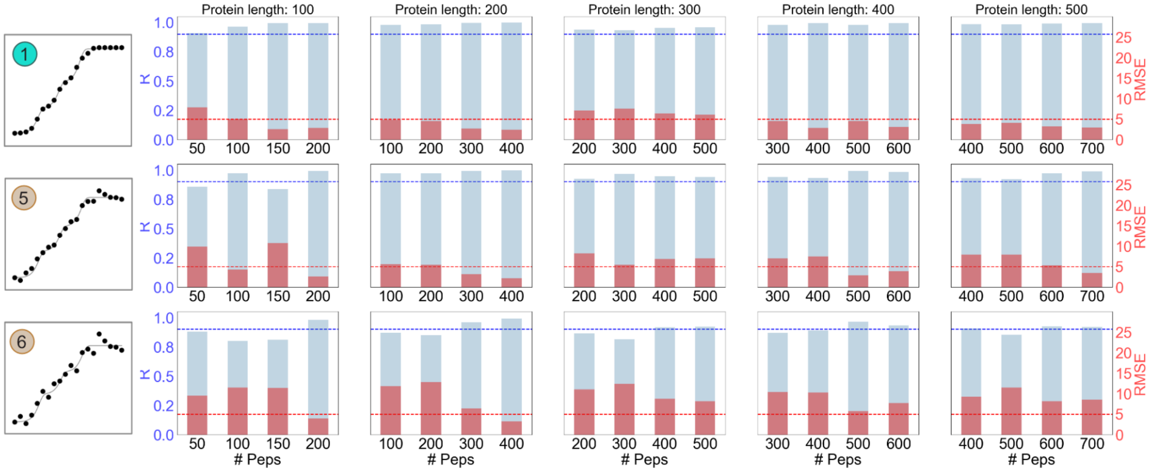

**Figure S2. FEATHER results with uncertainties < 1.0 log($k_{ex}$).**

FEATHER performance on synthetic datasets (perfect, perfect_noise10, and perfect_noise20 in Table S1), with R (blue) and RMSE (red). Only one protein was evaluated for each case due to the computational cost of running FEATHER on so many proteins.

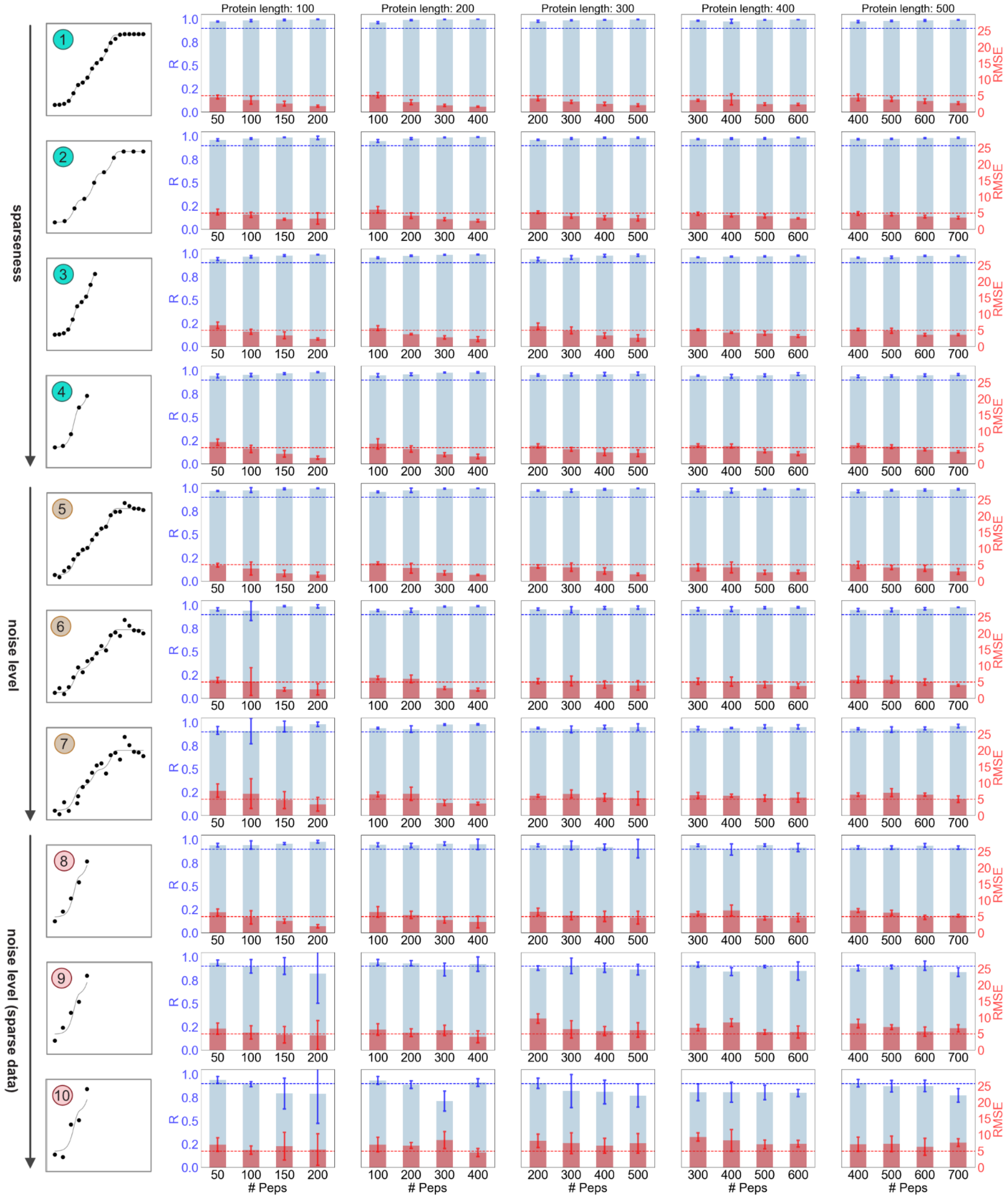

**Figure S3. Expanded data of Fig. 1b for PFNet-Centroid predictions with confidence > 0.8.**

PFNet-Centroid performance on synthetic datasets with varying degrees of sparsity and noise across proteins of different sizes and peptide numbers, with R (blue) and RMSE (red).

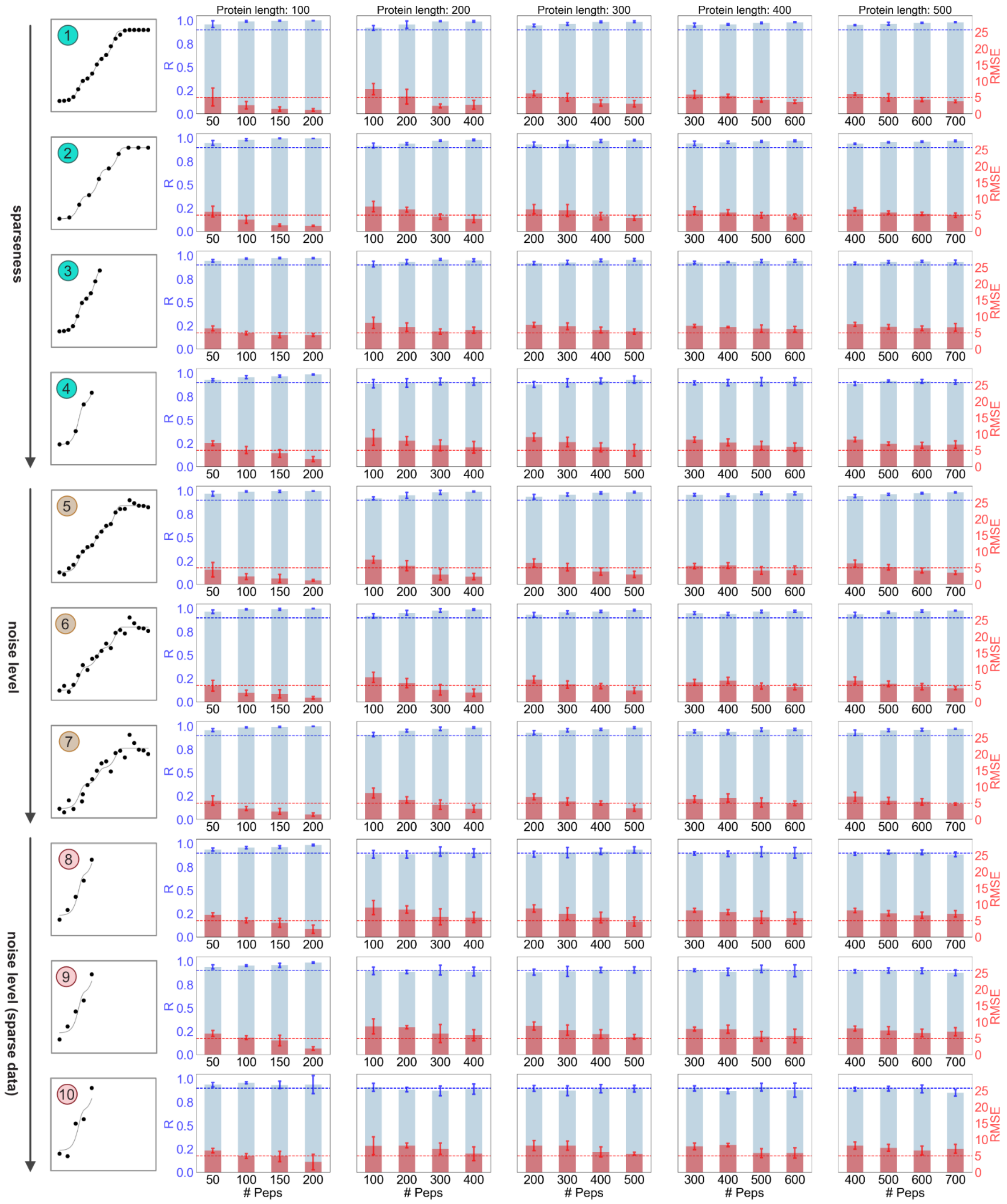

**Figure S4. Expanded data of Fig. 1b for PFNet(-CNN) predictions with confidence > 0.8.**

PFNet(-CNN) performance on synthetic datasets with varying degrees of sparsity and noise across proteins of different sizes and peptide numbers, with R (blue) and RMSE (red).

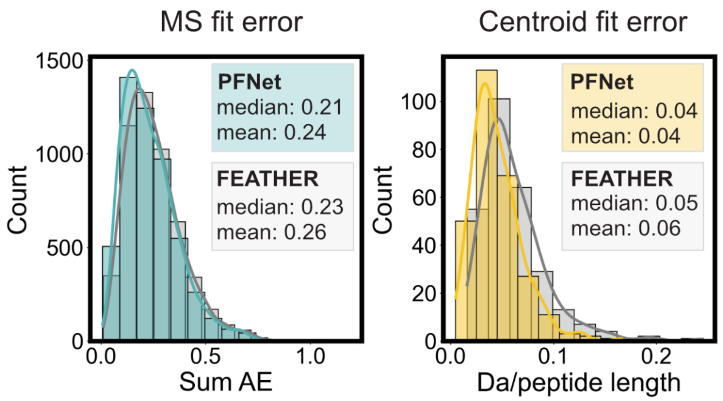

**Figure S5. Comparison of PFNet and FEATHER fitting errors for the mini-protein dataset.**

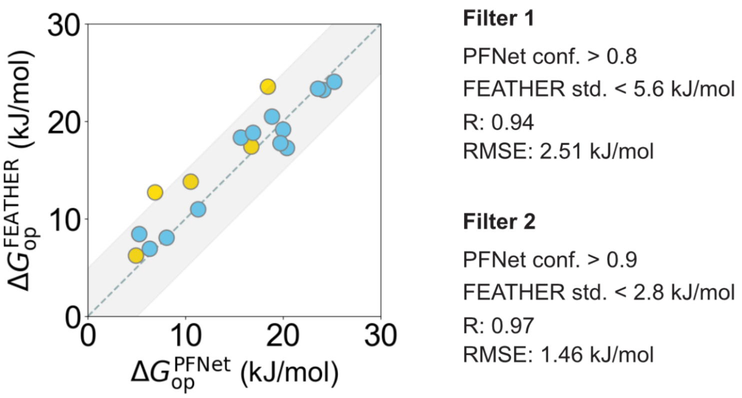

**Figure S6. Comparison of PFNet and FEATHER $\Delta G_{op}$ for the mini-protein dataset.**

R and RMSE were calculated at two different thresholds.

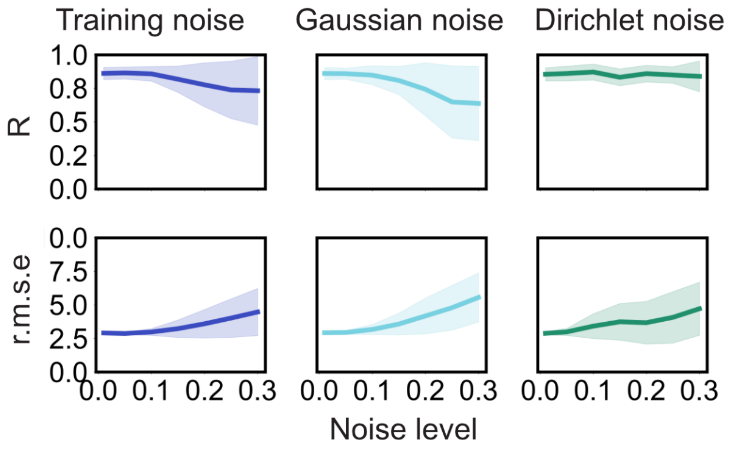

**Figure S7. PFNet robustness to additional noise injections in the mini-protein dataset.**

PFNet performance on the mini-protein dataset incorporating additive training, Gaussian, and Dirichlet noise. The shaded area indicates the variation across 20 independent runs.

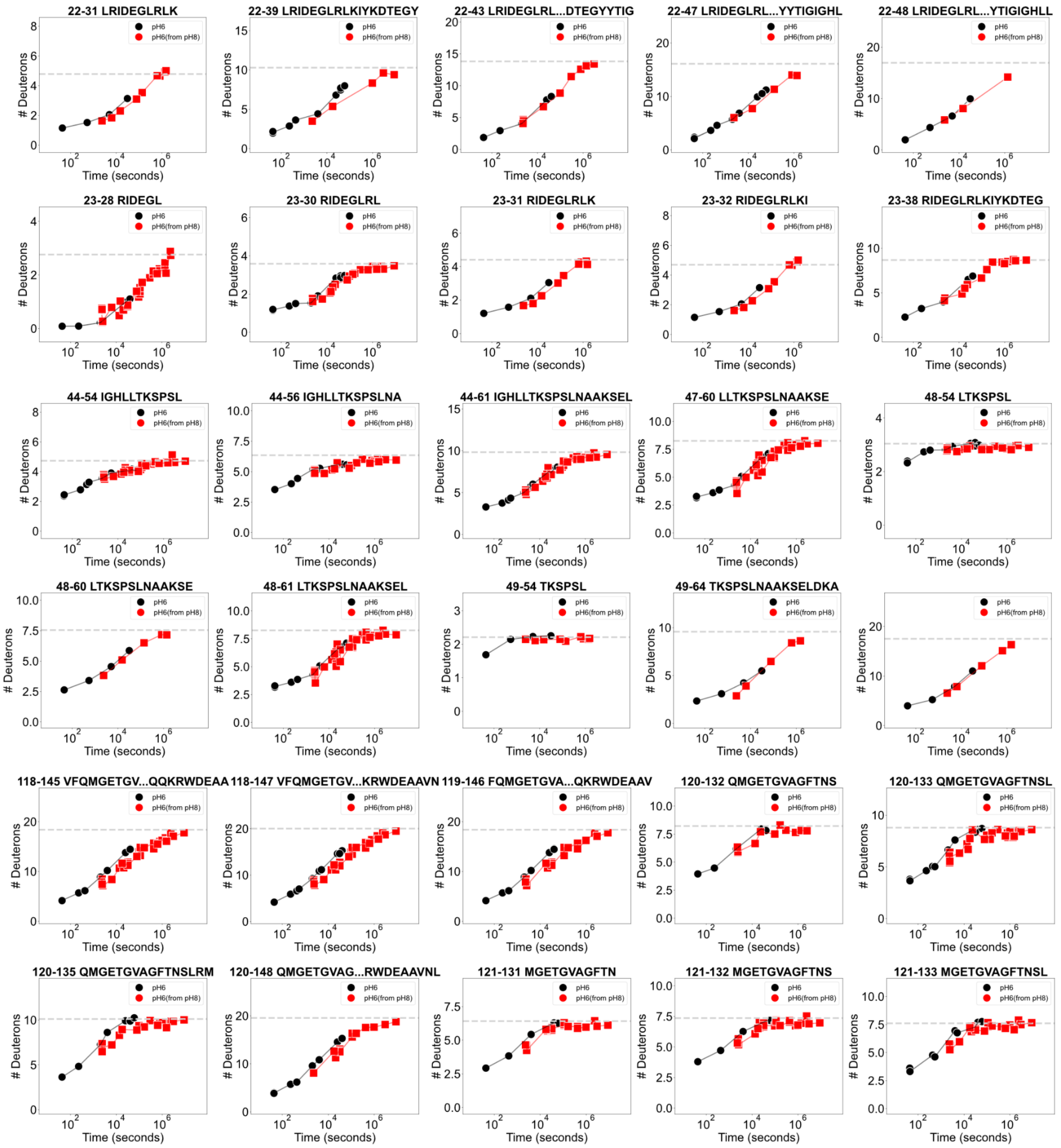

**Figure S8. Expanded exchange time window by pH conversion for T4L.**

Representative peptide uptake plots from HX/MS experiments comparing direct measurements at pH 6.0 with values converted from pH 8.0, illustrating the broadened exchange time coverage achieved through pH conversion.

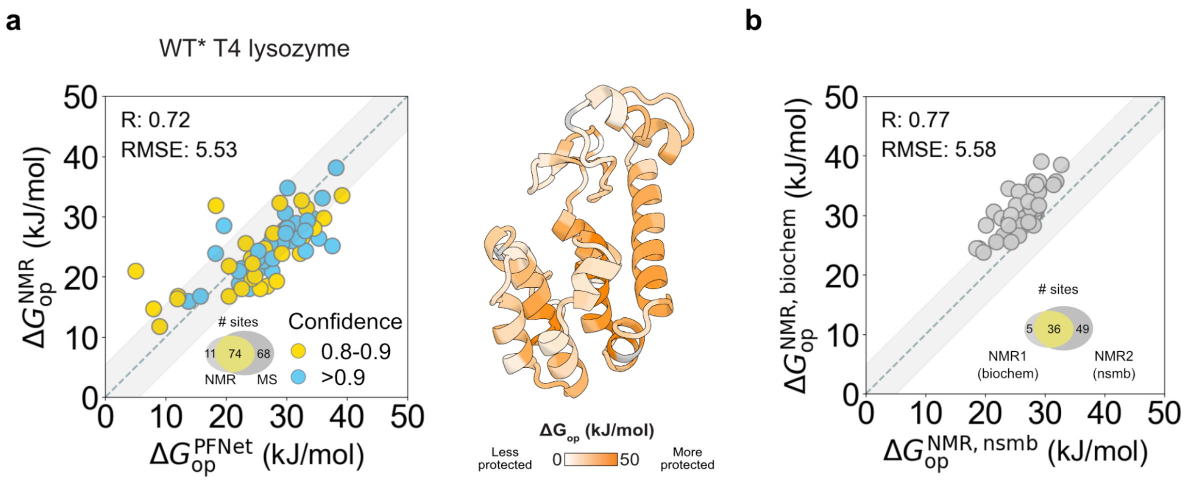

**Figure S9. Experimental benchmarking on the T4L dataset.**

**a,** PFNet-derived $\Delta G_{op}$ values mapped onto the crystal structure of T4L (PDB: 5VNR). **b,** Comparison of $\Delta G_{op}$ values obtained from two independent HX/NMR studies: NMR1 (*Biochemistry* 39, 248–254, 2000) and NMR2 (*Nature Structural Biology* 6, 1072–1078, 1999). The Venn diagram shows the number of single-residue sites resolved by each technique/studies.

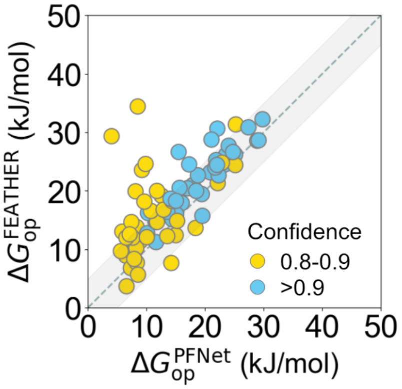

**Filter 1**

PFNet conf. > 0.8

FEATHER std. < 5.6 kJ/mol

R: 0.65

RMSE: 6.54 kJ/mol

**Filter 2**

PFNet conf. > 0.9

FEATHER std. < 2.8 kJ/mol

R: 0.87

RMSE: 3.58 kJ/mol

**Figure S10. Comparison of PFNet- and FEATHER-derived results on the T4L dataset.** R and RMSE were calculated at two different thresholds.

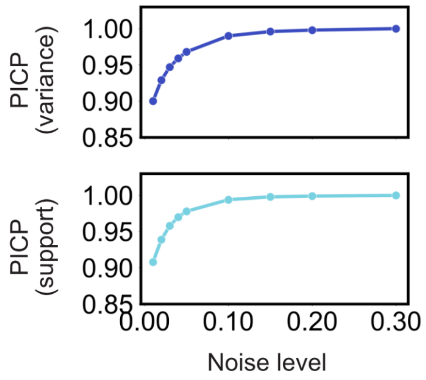

**Figure S11. PICP analysis for the T4L HX/MS dataset.**

The PICP analysis estimates the equivalent noise levels corresponding to the synthetic benchmarks.

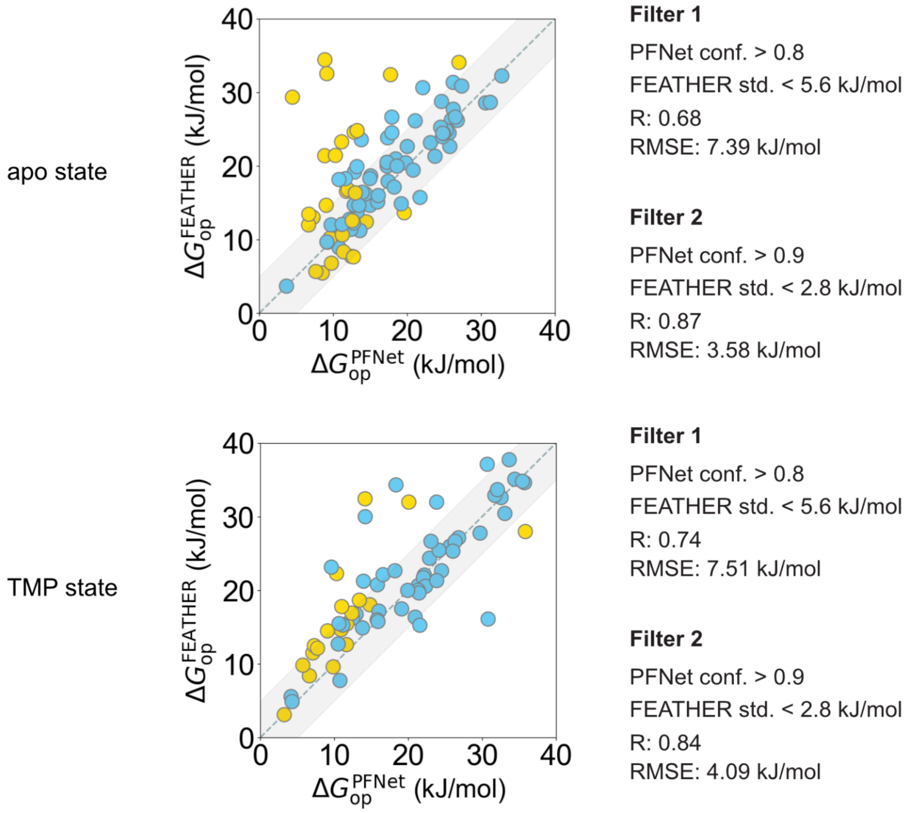

**Figure S12. Comparison of PFNet and FEATHER $\Delta G_{op}$ for apo and TMP-ecDHFR.**

R and RMSE were calculated at two different thresholds.

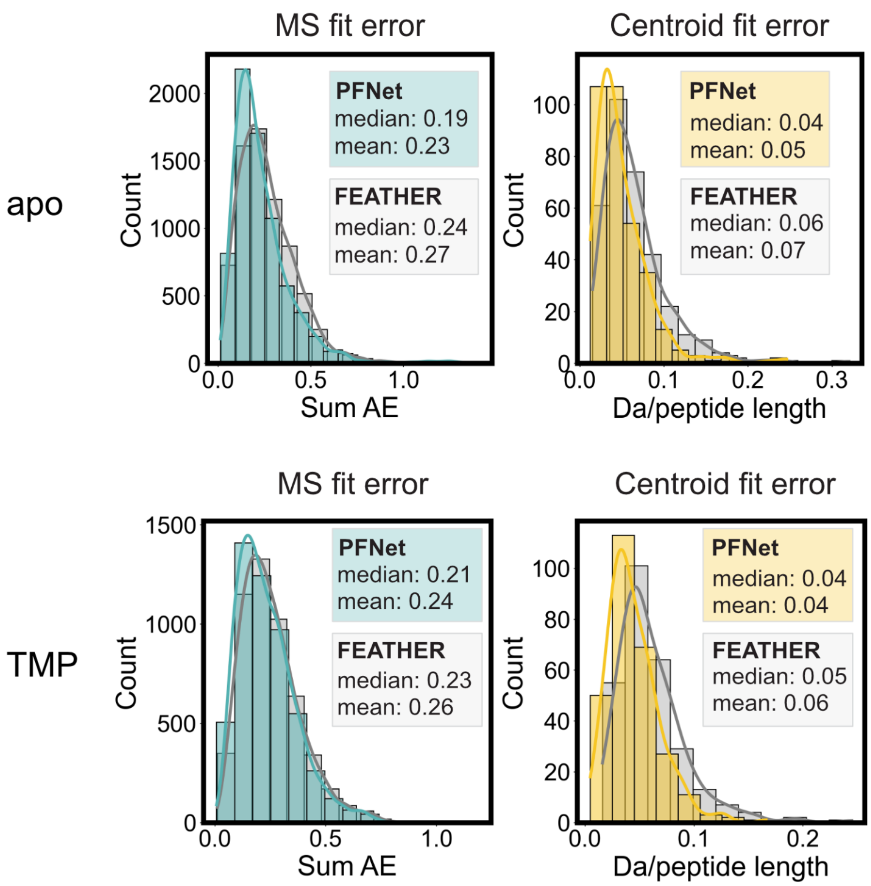

**Figure S13. Comparison of PFNet and FEATHER fitting errors on ecDHFR.**

An expanded version of Fig. 2b showing detailed statistical comparisons of fitting errors between PFNet and FEATHER.

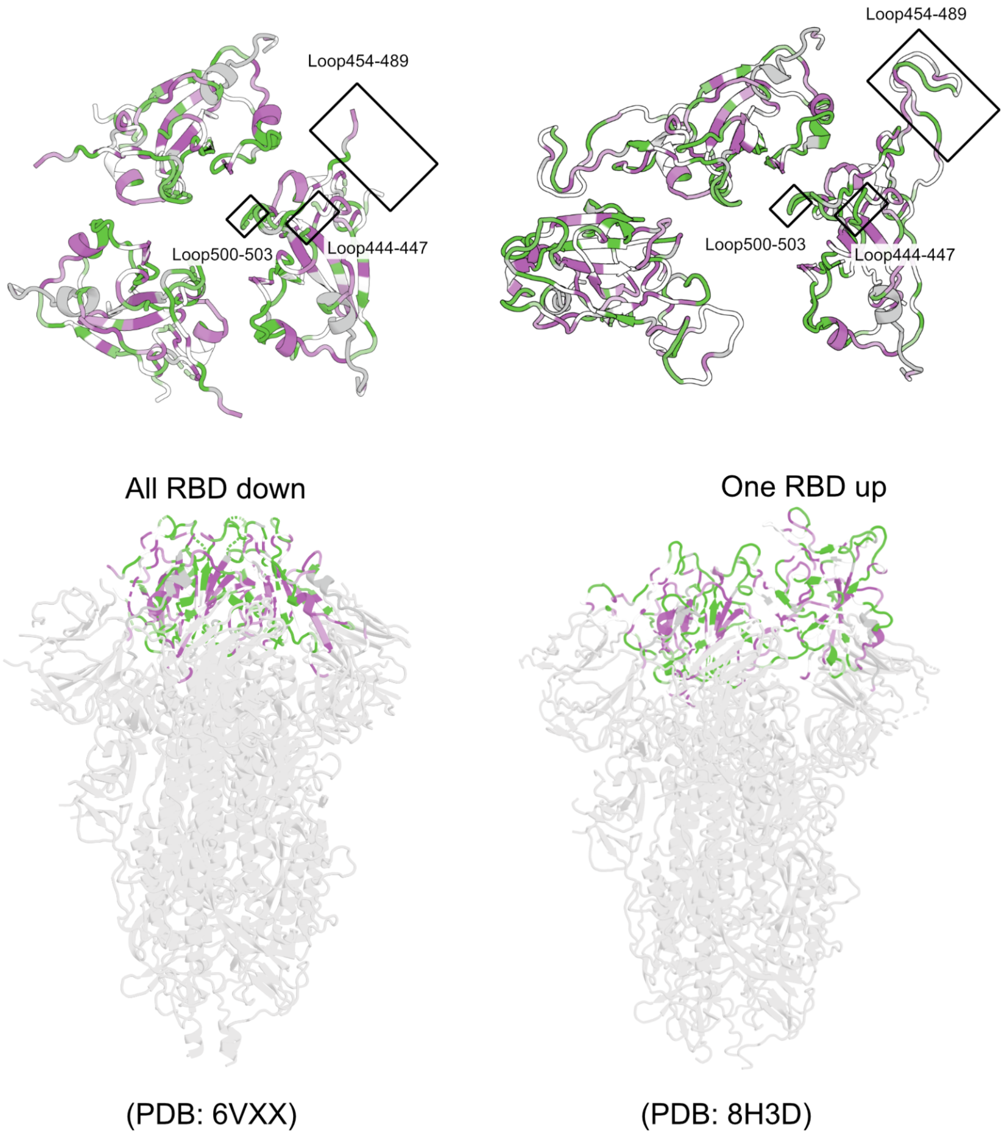

**Figure S14. PFNet-derived $\Delta\Delta G_{op(apo\text{-}ACE2)}$ values mapped on the spike structure.**

Three loop regions become rigidified upon the ACE2 binding-driven population shift in the spike protein, as shown in the all-RBD-down (PDB: 6VXX, left) and one RBD-up conformations (PDB: 8H3D, right).

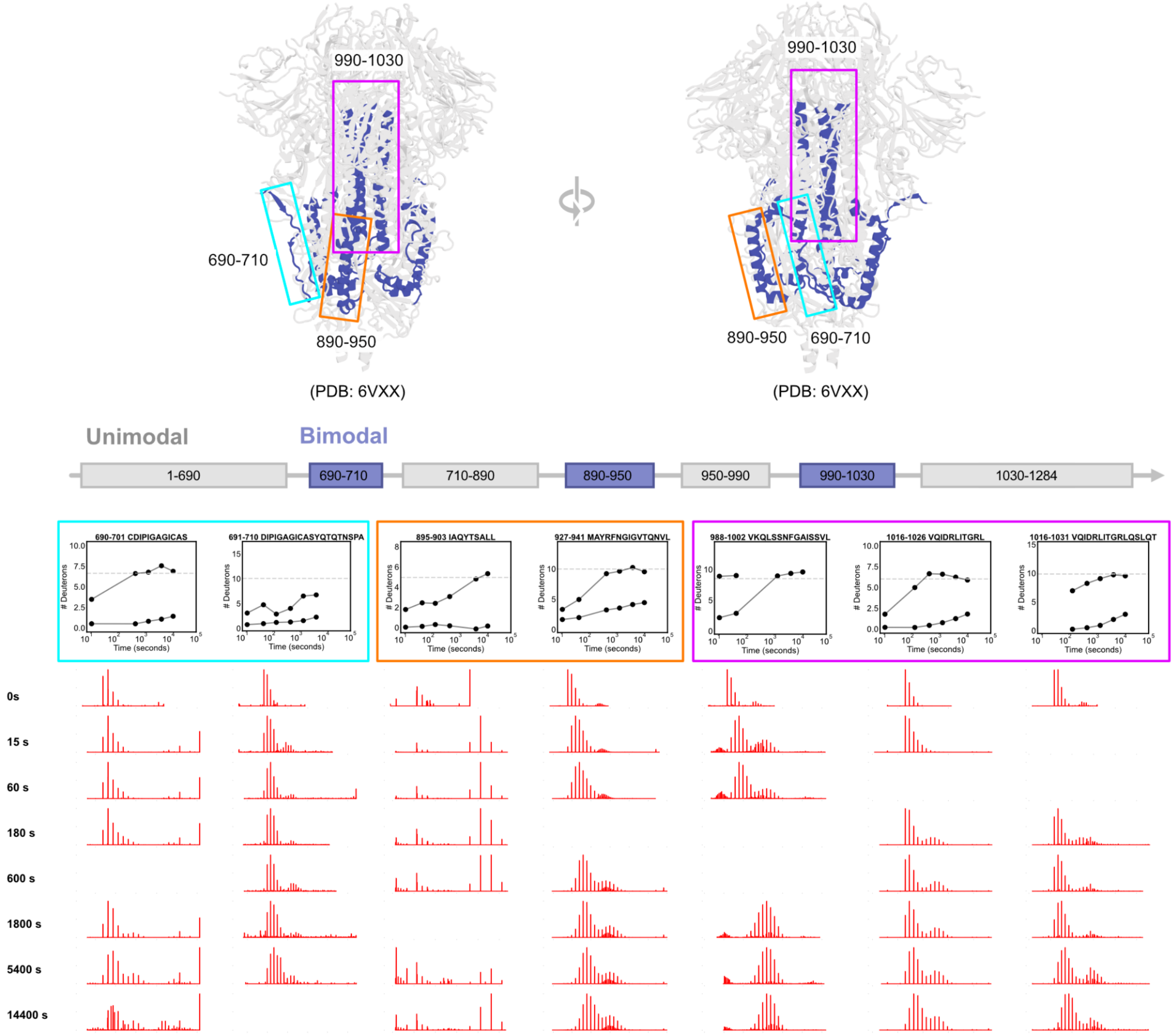

**Figure S15. Bimodal regions of the SARS-CoV-2 spike protein.**

Representative peptides shown for each region, alongside the raw isotopic mass envelopes.

### 2. Supplemental tables

#### Table S1. Synthetic benchmarking datasets

| Dataset index | Data quality | log(P) range | # timepoints | Time window (s) | Noise level |
|---|---|---|---|---|---|
| 1 | perfect | 2-12 | 20 | 1e-6 to 1e12 | 0 |
| 2 | sparse | 2-12 | 10 | 1e1 to 1e12 | 0 |
| 3 | tp_1e6 | 2-12 | 10 | 1e1 to 1e6 | 0 |
| 4 | sparse_tp_1e5 | 2-12 | 5 | 1e1 to 1e5 | 0 |
| 5 | perfect_noise10 | 2-12 | 20 | 1e1 to 1e12 | 0.1 |
| 6 | perfect_noise20 | 2-12 | 20 | 1e1 to 1e12 | 0.2 |
| 7 | perfect_noise30 | 2-12 | 20 | 1e1 to 1e12 | 0.3 |
| 8 | sparse_tp_1e5_noise10 | 2-12 | 5 | 1e1 to 1e5 | 0.1 |
| 9 | sparse_tp_1e5_noise20 | 2-12 | 5 | 1e1 to 1e5 | 0.2 |
| 10 | sparse_tp_1e5_noise30 | 2-12 | 5 | 1e1 to 1e5 | 0.3 |

**Table S2. HX experiment summary table for EEHEE_rd4_087.**

These experimental conditions were previously reported (Lu *et al.*, *Nature Chemical Biology*, 2025, https://doi.org/10.1038/s41589-025-02049-1) but are included here for convenience. All experiments were performed with the KEEP setting in the PIGEON peptide disambiguation software. Abbreviations: FP, immobilized protease type XIII. Nep, nepenthesin 2. AP, alanyl aminopeptidase.

| Dataset | **EEHEE_rd4_087 Rep1** | **EEHEE_rd4_087 Rep2** | **EEHEE_rd4_087 Rep3** |
|---|---|---|---|
| Protein states | APO | APO | APO |
| Date | 08/19/24 | 09/10/24 | 10/23/24 |
| Protease column | Nep/pepsin | AP/pepsin | FP/pepsin |
| Reaction details | 25mM MES, pH 6.0, temperature 20 °C, final $D_2O$ concentration 86.3% | | |
| Time course (s) | 4.2e1-6.0e4 | 4.6e1-3.0e4 | 4.4e1-2.5e4 |
| # of time points | 6 | 6 | 5 |
| Control samples | Maximally-labeled sample (full-D) | | |
| Back-exchange (mean / IQR) | 0.23/0.07 | 0.29/0.13 | 0.20/0.12 |
| # of peptides | 35 | 21 | 46 |
| Sequence coverage | 0.96 | 0.96 | 0.96 |
| Average peptide length / Redundancy | 1.3 | 2.2 | 1.0 |

**Table S3. $\Delta G_{op}$ values for EEHEE_rd4_087.**

| Res. name | Res. number | Apo | pStd Dev | Confidence | Single resolve |
|---|---|---|---|---|---|
| GLY | 1 | | | | |
| SER | 2 | 31.98 | 3.08 | 0.64 | FALSE |
| HIS | 3 | 25.56 | 3.08 | 0.63 | FALSE |
| MET | 4 | 24.04 | 3.08 | 0.89 | FALSE |
| THR | 5 | 20.09 | 3.08 | 0.92 | FALSE |
| GLN | 6 | 18.4 | 3.24 | 0.86 | FALSE |
| VAL | 7 | 14.14 | 3.24 | 0.91 | FALSE |
| HIS | 8 | 16.73 | 4.81 | 0.87 | TRUE |
| VAL | 9 | 11.03 | 3.99 | 0.9 | FALSE |
| ASP | 10 | 10.56 | 3.99 | 0.89 | FALSE |
| GLY | 11 | 14.6 | 12.37 | 0.7 | TRUE |
| VAL | 12 | 15.66 | 3.56 | 0.9 | TRUE |
| THR | 13 | 10.53 | 3.82 | 0.9 | TRUE |
| TYR | 14 | 19.99 | 2.27 | 0.94 | TRUE |
| THR | 15 | 8.06 | 2.1 | 0.94 | TRUE |
| PHE | 16 | 16.91 | 2.63 | 0.93 | TRUE |
| SER | 17 | 5.26 | 3.5 | 0.91 | TRUE |
| ASN | 18 | 18.43 | 4.4 | 0.88 | TRUE |
| PRO | 19 | | | | |
| GLU | 20 | 6.88 | 4.14 | 0.89 | TRUE |
| GLU | 21 | 6.34 | 1.8 | 0.95 | TRUE |
| ALA | 22 | 23.68 | 2.17 | 0.95 | FALSE |
| LYS | 23 | 24.17 | 2.17 | 0.94 | FALSE |
| LYS | 24 | 24.11 | 2.32 | 0.94 | TRUE |
| PHE | 25 | 18.85 | 2.12 | 0.94 | TRUE |
| ALA | 26 | 23.55 | 2.04 | 0.94 | TRUE |
| ASP | 27 | 25.24 | 2.05 | 0.95 | FALSE |
| GLU | 28 | 23.16 | 2.05 | 0.94 | FALSE |
| MET | 29 | 25.23 | 2.26 | 0.94 | TRUE |

| | | | | | |
|---|---|---|---|---|---|
| ALA | 30 | 20.38 | 2.65 | 0.93 | TRUE |
| LYS | 31 | 19.72 | 2.29 | 0.94 | TRUE |
| ARG | 32 | 13.43 | 4.03 | 0.94 | FALSE |
| LYS | 33 | 13.32 | 4.03 | 0.95 | FALSE |
| GLY | 34 | 12.81 | 4.03 | 0.94 | FALSE |
| GLY | 35 | 13.09 | 4.03 | 0.89 | FALSE |
| THR | 36 | 5.76 | 3.76 | 0.92 | FALSE |
| TRP | 37 | 4.02 | 3.76 | 0.9 | FALSE |
| GLU | 38 | 4.92 | 3.88 | 0.9 | TRUE |
| ILE | 39 | 3.86 | 3.96 | 0.76 | FALSE |
| LYS | 40 | 7.15 | 3.96 | 0.89 | FALSE |
| ASP | 41 | 11.29 | 3.21 | 0.91 | TRUE |
| GLY | 42 | 11.34 | 4.49 | 0.57 | FALSE |
| HIS | 43 | 17.39 | 4.49 | 0.88 | FALSE |
| ILE | 44 | 12.55 | 4.49 | 0.88 | FALSE |
| HIS | 45 | 15.15 | 4.49 | 0.92 | FALSE |
| VAL | 46 | 12.72 | 4.49 | 0.88 | FALSE |
| GLU | 47 | 2.52 | 6.07 | 0.84 | TRUE |

**Table S4. HX experiment summary table for T4 lysozyme.**

All experiments were performed with KEEP setting in PIGEON. Abbreviations: FP, immobilized protease type XIII. Nep, nepenthesin 2. AP, alanyl aminopeptidase.

| Dataset | **AP_pH8_Digest1** | **AP_pH8_Digest2** | **FP_pH8_Digest** |
|---|---|---|---|
| Protein states | APO | APO | APO |
| Date | 11/02/2024 | 03/18/25 | 03/14/25 |
| Protease column | AP/pepsin | AP/pepsin | FP/pepsin |
| Reaction details | 50 mM tris hydrochloride, 150 mM NaCl, 0.5 mM TCEP, pH 8.0, temperature 15 °C, final $D_2O$ concentration 90.9% | | |
| Time course (s) | 4.3e1-2.5e4 | 4.4e1-4.5e4 | 4.2e1-1.6e5 |
| # of time points | 5 | 4 | 6 |
| Control samples | Maximally-labeled sample (full-D) | | |
| Back-exchange (mean / IQR) | 0.23/0.08 | 0.25/0.08 | 0.23/0.08 |
| # of peptides | 168 | 182 | 215 |
| Sequence coverage | 0.92 | 0.93 | 0.93 |
| Average peptide length / Redundancy | 1.1 | 1.0 | 0.8 |

| | **N1__pH8_Digest** | **NP_pH8_Digest** | **AP_pH6_Digest** |
|---|---|---|---|
| Protein states | APO | APO | APO |
| Date | 03/24/25 | 11/04/24 | 05/10/25 |
| Protease column | Nepenthesin-1 | NP/pepsin | AP/pepsin |
| Reaction details | 50 mM tris hydrochloride, 150 mM NaCl, 0.5 mM TCEP, pH 8.0, temperature 15 °C, final $D_2O$ concentration 90.9% | | |
| Time course (s) | 4.3e1-2.5e4 | 4.4e1-2.8e4 | 4.3e1-4.0e4 |
| # of time points | 5 | 5 | 5 |
| Control samples | Maximally-labeled sample (full-D) | | |
| Back-exchange | 0.24/0.11 | 0.24/0.10 | 0.25/0.07 |

| | | | |
|---|---|---|---|
| (mean / IQR) | | | |
| # of peptides | 179 | 244 | 158 |
| Sequence coverage | 0.93 | 0.94 | 0.92 |
| Average peptide length / Redundancy | 1.0 | 0.7 | 1.1 |

| | **FP_pH6_Digest** | **N1_pH6_Digest** | **NP_pH6_Digest** |
|---|---|---|---|
| Protein states | APO | APO | APO |
| Date | 05/09/25 | 05/11/25 | 05/10/25 |
| Protease column | FP/pepsin | Nepenthesin-1 | NP/pepsin |
| Reaction details | 50 mM tris hydrochloride, 150 mM NaCl, 0.5 mM TCEP, pH 8.0, temperature 15 °C, final $D_2O$ concentration 90.9% | | |
| Time course (s) | 4.3e1-6.0e4 | 4.3e1-2.2e4 | 4.4e1-3.1e4 |
| # of time points | 5 | 4 | 4 |
| Control samples | Maximally-labeled sample (full-D) | | |
| Back-exchange (mean / IQR) | 0.23/0.08 | 0.24/0.11 | 0.23/0.11 |
| # of peptides | 199 | 130 | 190 |
| Sequence coverage | 0.93 | 0.93 | 0.94 |
| Average peptide length / Redundancy | 0.9 | 1.4 | 0.9 |

**Table S5. $\Delta G_{op}$ values for T4 lysozyme.**

| Res. name | Res. number | Apo | pStd Dev | Confidence | Single resolve |
|---|---|---|---|---|---|
| GLY | 1 | | | | |
| SER | 2 | 31.98 | 3.08 | 0.64 | FALSE |
| HIS | 3 | 25.56 | 3.08 | 0.63 | FALSE |
| MET | 4 | 24.04 | 3.08 | 0.89 | FALSE |
| THR | 5 | 20.09 | 3.08 | 0.92 | FALSE |
| GLN | 6 | 18.4 | 3.24 | 0.86 | FALSE |
| VAL | 7 | 14.14 | 3.24 | 0.91 | FALSE |
| HIS | 8 | 16.73 | 4.81 | 0.87 | TRUE |
| VAL | 9 | 11.03 | 3.99 | 0.9 | FALSE |
| ASP | 10 | 10.56 | 3.99 | 0.89 | FALSE |
| GLY | 11 | 14.6 | 12.37 | 0.7 | TRUE |
| VAL | 12 | 15.66 | 3.56 | 0.9 | TRUE |
| THR | 13 | 10.53 | 3.82 | 0.9 | TRUE |
| TYR | 14 | 19.99 | 2.27 | 0.94 | TRUE |
| THR | 15 | 8.06 | 2.1 | 0.94 | TRUE |
| PHE | 16 | 16.91 | 2.63 | 0.93 | TRUE |
| SER | 17 | 5.26 | 3.5 | 0.91 | TRUE |
| ASN | 18 | 18.43 | 4.4 | 0.88 | TRUE |
| PRO | 19 | | | | |
| GLU | 20 | 6.88 | 4.14 | 0.89 | TRUE |
| GLU | 21 | 6.34 | 1.8 | 0.95 | TRUE |
| ALA | 22 | 23.68 | 2.17 | 0.95 | FALSE |
| LYS | 23 | 24.17 | 2.17 | 0.94 | FALSE |
| LYS | 24 | 24.11 | 2.32 | 0.94 | TRUE |
| PHE | 25 | 18.85 | 2.12 | 0.94 | TRUE |
| ALA | 26 | 23.55 | 2.04 | 0.94 | TRUE |
| ASP | 27 | 25.24 | 2.05 | 0.95 | FALSE |
| GLU | 28 | 23.16 | 2.05 | 0.94 | FALSE |
| MET | 29 | 25.23 | 2.26 | 0.94 | TRUE |

| | | | | | |
|---|---|---|---|---|---|
| ALA | 30 | 20.38 | 2.65 | 0.93 | TRUE |
| LYS | 31 | 19.72 | 2.29 | 0.94 | TRUE |
| ARG | 32 | 13.43 | 4.03 | 0.94 | FALSE |
| LYS | 33 | 13.32 | 4.03 | 0.95 | FALSE |
| GLY | 34 | 12.81 | 4.03 | 0.94 | FALSE |
| GLY | 35 | 13.09 | 4.03 | 0.89 | FALSE |
| THR | 36 | 5.76 | 3.76 | 0.92 | FALSE |
| TRP | 37 | 4.02 | 3.76 | 0.9 | FALSE |
| GLU | 38 | 4.92 | 3.88 | 0.9 | TRUE |
| ILE | 39 | 3.86 | 3.96 | 0.76 | FALSE |
| LYS | 40 | 7.15 | 3.96 | 0.89 | FALSE |
| ASP | 41 | 11.29 | 3.21 | 0.91 | TRUE |
| GLY | 42 | 11.34 | 4.49 | 0.57 | FALSE |
| HIS | 43 | 17.39 | 4.49 | 0.88 | FALSE |
| ILE | 44 | 12.55 | 4.49 | 0.88 | FALSE |
| HIS | 45 | 15.15 | 4.49 | 0.92 | FALSE |
| VAL | 46 | 12.72 | 4.49 | 0.88 | FALSE |
| GLU | 47 | 2.52 | 6.07 | 0.84 | TRUE |

**Table S6. HX experiment summary table for SARS-CoV-2 spike protein.**

Data were obtained from Costello *et al.* (Nat. Struct. Mol. Biol. 2022, 29 (3), 229–238). The peptide list was generated using PIGEON (Lu *et al.*, *Nature Chemical Biology*, 2025, https://doi.org/10.1038/s41589-025-02049-1).

| Dataset | **Spike_APO_rep1** | **Spike_APO_rep2** | **Spike_APO_rep3** | **Spike_APO_rep4** |
|---|---|---|---|---|
| Protein states | APO | APO | APO | APO |
| Date | 07/09/20 | 07/13/20 | 07/14/20 | 03/15/21 |
| Time course (s) | 15-14400 | | | |
| # of time points | 7 | | | |
| Control samples | Maximally-labeled sample (full-D) | | | |
| Back-exchange (mean / IQR) | 0.39/0.14 | 0.38/0.13 | 0.38/0.13 | 0.39/0.14 |
| # of peptides | 887 | 701 | 701 | 801 |
| Sequence coverage | 0.95 | 0.92 | 0.93 | 0.95 |
| Average peptide length / Redundancy | 1.4 | 1.8 | 1.8 | 1.6 |

| Dataset | **Spike_ACE2_rep1** | **Spike_ACE2_rep2** | **Spike_ACE2_rep3** |
|---|---|---|---|
| Protein states | ACE2 | ACE2 | ACE2 |
| Date | 07/08/20 | 08/06/20 | 08/06/20 |
| Time course (s) | 15-14400 | | |
| # of time points | 7 | | |
| Control samples | Maximally-labeled sample (full-D) | | |
| Back-exchange (mean / IQR) | 0.40/0.14 | 0.40/0.14 | 0.40/0.14 |
| # of peptides | 572 | 684 | 688 |
| Sequence coverage | 0.95 | 0.98 | 0.98 |
| Average peptide length / Redundancy | 2.2 | 1.9 | 1.9 |

**Table S7. $\Delta G_{op}$ values for the RBD region of the SARS-CoV-2 spike protein**

| Res. name | Res. number | Apo | pStd Dev | Confidence | Single resolved | ACE2 | pStd Dev | Confidence | Single resolved |
|---|---|---|---|---|---|---|---|---|---|
| ALA | 250 | 39.9 | 12.09 | 0.87 | FALSE | 36.59 | 16.82 | 0.62 | TRUE |
| LEU | 251 | 34.88 | 12.09 | 0.78 | FALSE | 25.76 | 7.24 | 0.69 | FALSE |
| GLU | 252 | 36.38 | 12.09 | 0.53 | FALSE | 27.25 | 7.24 | 0.9 | FALSE |
| PRO | 253 | 0 | 0 | -100 | FALSE | 0 | 0 | -100 | FALSE |
| LEU | 254 | 23.69 | 2.4 | 0.94 | TRUE | 24.39 | 7.24 | 0.87 | FALSE |
| VAL | 255 | 25.63 | 2.87 | 0.92 | TRUE | 37.8 | 15.36 | 0.65 | TRUE |
| ASP | 256 | 4.75 | 3.99 | 0.89 | TRUE | 33.68 | 11.37 | 0.68 | FALSE |
| LEU | 257 | 40.94 | 16.6 | 0.63 | TRUE | 29.57 | 11.37 | 0.77 | FALSE |
| PRO | 258 | 0 | 0 | -100 | TRUE | 0 | 0 | -100 | FALSE |
| ILE | 259 | 7.87 | 10.72 | 0.74 | TRUE | 2.8 | 5.77 | 0.85 | TRUE |
| GLY | 260 | | | | | | | | |
| ILE | 261 | | | | | | | | |
| ASN | 262 | | | | | | | | |
| ILE | 263 | | | | | | | | |
| THR | 264 | | | | | | | | |
| ARG | 265 | | | | | | | | |
| PHE | 266 | | | | | | | | |
| GLN | 267 | | | | | | | | |
| THR | 268 | | | | | | | | |
| LEU | 269 | | | | | 34.35 | 16.08 | 0.63 | FALSE |
| LEU | 270 | | | | | 32.01 | 16.08 | 0.65 | FALSE |
| ALA | 271 | 27.56 | 17.47 | 0.61 | TRUE | 47.05 | 17.47 | 0.61 | TRUE |
| LEU | 272 | 10.36 | 10.64 | 0.62 | FALSE | 15.84 | 8.63 | 0.66 | FALSE |
| HIS | 273 | 14.77 | 10.64 | 0.88 | FALSE | 20.25 | 8.63 | 0.64 | FALSE |
| ARG | 274 | 17.55 | 9.74 | 0.56 | FALSE | 22.39 | 8.63 | 0.88 | FALSE |
| SER | 275 | 17.67 | 9.74 | 0.87 | FALSE | 22.51 | 8.63 | 0.9 | FALSE |
| TYR | 276 | 14.48 | 9.74 | 0.91 | FALSE | 19.32 | 8.63 | 0.91 | FALSE |
| LEU | 277 | 5.22 | 4.92 | 0.87 | TRUE | 5.14 | 5.45 | 0.86 | TRUE |
| THR | 278 | 5.26 | 4.99 | 0.82 | FALSE | 9.93 | 6.95 | 0.71 | FALSE |

| | | | | | | | | | |
|---|---|---|---|---|---|---|---|---|---|
| PRO | 279 | | | | | | | | |
| GLY | 280 | 5.32 | 4.99 | 0.92 | FALSE | 9.99 | 6.95 | 0.8 | FALSE |
| ASP | 281 | 17.14 | 10.69 | 0.61 | FALSE | 13.08 | 6.95 | 0.89 | FALSE |
| SER | 282 | 16.7 | 10.69 | 0.75 | FALSE | 12.63 | 6.95 | 0.91 | FALSE |
| SER | 283 | 19.42 | 10.69 | 0.89 | FALSE | 21.1 | 13.42 | 0.69 | TRUE |
| SER | 284 | 11.11 | 5.29 | 0.76 | FALSE | 10.88 | 3.46 | 0.89 | FALSE |
| GLY | 285 | 8.83 | 5.29 | 0.92 | FALSE | 8.59 | 3.46 | 0.91 | FALSE |
| TRP | 286 | 5.92 | 5.29 | 0.92 | FALSE | 5.68 | 3.46 | 0.91 | FALSE |
| THR | 287 | 7.59 | 4.01 | 0.93 | FALSE | 6.02 | 3.46 | 0.92 | FALSE |
| ALA | 288 | 9.76 | 4.01 | 0.88 | FALSE | 11.37 | 5.55 | 0.68 | FALSE |
| GLY | 289 | 8.45 | 4.01 | 0.88 | FALSE | 10.06 | 5.55 | 0.89 | FALSE |
| ALA | 290 | 15.27 | 6.14 | 0.84 | TRUE | 11.2 | 5.55 | 0.91 | FALSE |
| ALA | 291 | 8.08 | 3.72 | 0.9 | FALSE | 10.23 | 5.55 | 0.92 | FALSE |
| ALA | 292 | 8.08 | 3.72 | 0.91 | FALSE | 10.23 | 5.55 | 0.91 | FALSE |
| TYR | 293 | 5.93 | 3.69 | 0.9 | TRUE | 24.61 | 3.63 | 0.9 | TRUE |
| TYR | 294 | 4.77 | 3.29 | 0.91 | TRUE | 20.1 | 4.17 | 0.89 | TRUE |
| VAL | 295 | 36.19 | 21.38 | 0.55 | TRUE | 31.68 | 7.64 | 0.74 | FALSE |
| GLY | 296 | 22.3 | 3.48 | 0.91 | TRUE | 34.42 | 7.64 | 0.77 | FALSE |
| TYR | 297 | 52.3 | 9.79 | 0.76 | TRUE | 34.82 | 7.64 | 0.93 | FALSE |
| LEU | 298 | 37.12 | 8.29 | 0.79 | TRUE | 41.69 | 19.4 | 0.58 | TRUE |
| GLN | 299 | 52.95 | 9.01 | 0.78 | TRUE | 46.18 | 14.36 | 0.67 | TRUE |
| PRO | 300 | | | | | | | | |
| ARG | 301 | 52.29 | 9.62 | 0.76 | TRUE | 51.52 | 8.3 | 0.79 | FALSE |
| THR | 302 | 14.2 | 1.64 | 0.96 | TRUE | 12.92 | 5.05 | 0.87 | TRUE |
| PHE | 303 | 16.19 | 8.4 | 0.79 | TRUE | 32.26 | 15.04 | 0.66 | TRUE |
| LEU | 304 | 49.98 | 12.2 | 0.71 | TRUE | 50.95 | 10.99 | 0.74 | TRUE |
| LEU | 305 | 31.29 | 23.23 | 0.52 | TRUE | 41.81 | 12.18 | 0.71 | TRUE |
| LYS | 306 | 16.7 | 10.17 | 0.59 | FALSE | 3.88 | 4.24 | 0.89 | TRUE |
| TYR | 307 | 17.27 | 10.17 | 0.71 | FALSE | 5.3 | 4.38 | 0.85 | FALSE |
| ASN | 308 | 21.2 | 10.17 | 0.85 | FALSE | 9.24 | 4.38 | 0.89 | FALSE |
| GLU | 309 | 19.33 | 10.17 | 0.89 | FALSE | 7.36 | 4.38 | 0.91 | FALSE |

| | | | | | | | | | |
|---|---|---|---|---|---|---|---|---|---|
| ASN | 310 | 32.39 | 17.17 | 0.59 | FALSE | 14.27 | 4.42 | 0.89 | FALSE |
| GLY | 311 | 32.09 | 17.17 | 0.65 | FALSE | 13.97 | 4.42 | 0.88 | FALSE |
| THR | 312 | 22.13 | 7.56 | 0.62 | FALSE | 24.5 | 4.76 | 0.93 | FALSE |
| ILE | 313 | 18.53 | 7.56 | 0.69 | FALSE | 20.91 | 4.76 | 0.82 | FALSE |
| THR | 314 | 19.84 | 7.56 | 0.85 | FALSE | 18.97 | 10.97 | 0.59 | FALSE |
| ASP | 315 | 23.28 | 7.56 | 0.87 | FALSE | 22.41 | 10.97 | 0.91 | FALSE |
| ALA | 316 | 20.54 | 7.56 | 0.87 | FALSE | 31.78 | 15.75 | 0.64 | TRUE |
| VAL | 317 | 17.56 | 7.56 | 0.88 | FALSE | 50.83 | 10.87 | 0.74 | TRUE |
| ASP | 318 | 21.34 | 7.56 | 0.88 | FALSE | 36.94 | 15.51 | 0.75 | FALSE |
| CYS | 319 | 24.08 | 7.56 | 0.87 | FALSE | 39.69 | 15.51 | 0.56 | FALSE |
| ALA | 320 | 6.43 | 4.47 | 0.88 | TRUE | 27.28 | 23.79 | 0.51 | TRUE |
| LEU | 321 | 45.46 | 14.24 | 0.67 | TRUE | 44.39 | 12.36 | 0.71 | TRUE |
| ASP | 322 | 37.43 | 7.14 | 0.82 | TRUE | 28.63 | 4.51 | 0.88 | TRUE |
| PRO | 323 | | | | | | | | |
| LEU | 324 | 14.36 | 8.03 | 0.8 | TRUE | 38.53 | 12.25 | 0.71 | FALSE |
| SER | 325 | 27.03 | 4.02 | 0.88 | FALSE | 4.88 | 3.66 | 0.9 | TRUE |
| GLU | 326 | 27.21 | 4.02 | 0.89 | FALSE | 25.5 | 4.97 | 0.87 | TRUE |
| THR | 327 | 24.89 | 4.02 | 0.89 | FALSE | 20.04 | 3.88 | 0.9 | TRUE |
| LYS | 328 | 27.03 | 4.02 | 0.91 | FALSE | 14.4 | 6.54 | 0.83 | TRUE |
| CYS | 329 | 46.11 | 13.81 | 0.68 | TRUE | 22.91 | 3.33 | 0.91 | TRUE |
| THR | 330 | 42.92 | 19.8 | 0.57 | TRUE | 37.99 | 9.01 | 0.85 | FALSE |
| LEU | 331 | 51.13 | 11.18 | 0.73 | TRUE | 33.08 | 9.01 | 0.71 | FALSE |
| LYS | 332 | 11.78 | 7.08 | 0.82 | TRUE | 19.45 | 5.02 | 0.87 | TRUE |
| SER | 333 | 10.11 | 5.58 | 0.86 | TRUE | 53.54 | 9.07 | 0.8 | FALSE |
| PHE | 334 | 22.84 | 7.2 | 0.82 | TRUE | 51.08 | 9.07 | 0.75 | FALSE |
| THR | 335 | 11.63 | 5.27 | 0.86 | TRUE | 11.67 | 10.82 | 0.74 | TRUE |
| VAL | 336 | 2.23 | 2.81 | 0.92 | FALSE | 2.07 | 2.71 | 0.93 | FALSE |
| GLU | 337 | 3.67 | 2.81 | 0.93 | FALSE | 3.51 | 2.71 | 0.93 | FALSE |
| LYS | 338 | 4.02 | 2.81 | 0.92 | FALSE | 3.86 | 2.71 | 0.92 | FALSE |
| GLY | 339 | 26.88 | 12.29 | 0.71 | TRUE | 52.8 | 9.15 | 0.77 | TRUE |
| ILE | 340 | 21.93 | 2.69 | 0.93 | TRUE | 22.99 | 2.61 | 0.93 | TRUE |

| | | | | | | | | | |
|---|---|---|---|---|---|---|---|---|---|
| TYR | 341 | 20.34 | 3.29 | 0.91 | TRUE | 12.02 | 7.51 | 0.81 | TRUE |
| GLN | 342 | 9.04 | 5.1 | 0.87 | TRUE | 6.05 | 3.88 | 0.9 | TRUE |
| THR | 343 | 8.68 | 4.84 | 0.87 | TRUE | 6.96 | 5.08 | 0.87 | TRUE |
| SER | 344 | 8.86 | 4.95 | 0.87 | TRUE | 15.21 | 3.34 | 0.93 | FALSE |
| ASN | 345 | 6.4 | 3.94 | 0.9 | TRUE | 16.47 | 3.34 | 0.89 | FALSE |
| PHE | 346 | 15.81 | 5.29 | 0.86 | TRUE | 18.99 | 5.14 | 0.87 | TRUE |
| ARG | 347 | 6.95 | 3.67 | 0.9 | TRUE | 18.46 | 4.26 | 0.89 | TRUE |
| VAL | 348 | 9.24 | 2.42 | 0.94 | FALSE | 5.31 | 2.81 | 0.92 | TRUE |
| GLN | 349 | 11.53 | 2.42 | 0.93 | FALSE | 18.3 | 2.91 | 0.92 | FALSE |
| PRO | 350 | | | | | | | | |
| THR | 351 | 3.18 | 3.02 | 0.92 | FALSE | 5.26 | 4.02 | 0.89 | TRUE |
| GLU | 352 | 5.48 | 3.02 | 0.92 | FALSE | 7.44 | 4.78 | 0.88 | TRUE |
| SER | 353 | 6.23 | 3.02 | 0.91 | FALSE | 3.16 | 3.49 | 0.91 | TRUE |
| ILE | 354 | 30.64 | 8.47 | 0.79 | TRUE | 7.89 | 4.86 | 0.88 | FALSE |
| VAL | 355 | 5.23 | 2.68 | 0.93 | TRUE | 5.03 | 4.86 | 0.86 | FALSE |
| ARG | 356 | 2.66 | 3.26 | 0.91 | TRUE | 12 | 7.54 | 0.81 | TRUE |
| PHE | 357 | 8.97 | 4.61 | 0.88 | TRUE | 10.36 | 3.92 | 0.92 | FALSE |
| PRO | 358 | | | | | | | | |
| ASN | 359 | 13.58 | 2.96 | 0.93 | FALSE | 11.9 | 3.92 | 0.88 | FALSE |
| ILE | 360 | 9.81 | 2.96 | 0.91 | FALSE | 3.25 | 3.49 | 0.91 | TRUE |
| THR | 361 | | | | | 29.67 | 5.49 | 0.86 | TRUE |
| ASN | 362 | | | | | 30.19 | 7.83 | 0.85 | FALSE |
| LEU | 363 | | | | | 24.77 | 7.83 | 0.76 | FALSE |
| CYS | 364 | | | | | 7.91 | 3.94 | 0.9 | FALSE |
| PRO | 365 | | | | | 0 | 0 | | |
| PHE | 366 | | | | | 13.63 | 6.54 | 0.83 | TRUE |
| GLY | 367 | | | | | 10.33 | 4.09 | 0.89 | TRUE |
| GLU | 368 | | | | | 10.62 | 3.25 | 0.89 | FALSE |
| VAL | 369 | | | | | 5.44 | 3.25 | 0.94 | FALSE |
| PHE | 370 | | | | | 46.14 | 7.56 | 0.8 | FALSE |
| ASN | 371 | 36.52 | 13.44 | 0.58 | FALSE | 51.45 | 7.56 | 0.82 | FALSE |

| | | | | | | | | | |
|---|---|---|---|---|---|---|---|---|---|
| ALA | 372 | 35.21 | 13.44 | 0.69 | FALSE | 27.73 | 3.79 | 0.9 | TRUE |
| THR | 373 | 32.98 | 13.44 | 0.65 | FALSE | 24.73 | 4 | 0.89 | TRUE |
| ARG | 374 | 34.98 | 13.44 | 0.84 | FALSE | 16.85 | 8.63 | 0.79 | TRUE |
| PHE | 375 | 37.41 | 12.77 | 0.69 | FALSE | 16.86 | 7.16 | 0.82 | TRUE |
| ALA | 376 | 37.87 | 12.77 | 0.71 | FALSE | 52 | 10.61 | 0.74 | TRUE |
| SER | 377 | 49.17 | 11.27 | 0.76 | FALSE | 43.93 | 9.82 | 0.8 | FALSE |
| VAL | 378 | 44.77 | 11.27 | 0.7 | FALSE | 39.53 | 9.82 | 0.8 | FALSE |
| TYR | 379 | 24.88 | 26.95 | 0.47 | TRUE | 39.47 | 9.82 | 0.69 | FALSE |
| ALA | 380 | 53.39 | 8.63 | 0.79 | TRUE | 50.66 | 12.46 | 0.71 | TRUE |
| TRP | 381 | 31.51 | 7.27 | 0.72 | FALSE | 41.76 | 11.5 | 0.72 | FALSE |
| ASN | 382 | 36.02 | 7.27 | 0.92 | FALSE | 46.27 | 11.5 | 0.73 | FALSE |
| ARG | 383 | 51.54 | 11.66 | 0.72 | TRUE | 35.12 | 19.22 | 0.58 | TRUE |
| LYS | 384 | 4.89 | 2.88 | 0.92 | FALSE | 8.64 | 6.08 | 0.84 | TRUE |
| ARG | 385 | 5 | 2.88 | 0.93 | FALSE | 20.46 | 15.02 | 0.6 | FALSE |
| ILE | 386 | 11.42 | 12.18 | 0.71 | TRUE | 16.41 | 15.02 | 0.71 | FALSE |
| SER | 387 | 15.69 | 12.47 | 0.71 | TRUE | 35.91 | 12.4 | 0.68 | FALSE |
| ASN | 388 | 32.49 | 11.29 | 0.7 | FALSE | 39.62 | 12.4 | 0.77 | FALSE |
| CYS | 389 | 33.34 | 11.29 | 0.77 | FALSE | 40.47 | 12.4 | 0.68 | FALSE |
| VAL | 390 | 50.68 | 10.95 | 0.74 | TRUE | 31.77 | 14.84 | 0.66 | TRUE |
| ALA | 391 | 45.15 | 9.33 | 0.77 | FALSE | 38.47 | 18.21 | 0.6 | TRUE |
| ASP | 392 | 46.53 | 9.33 | 0.77 | FALSE | 36.62 | 22.4 | 0.54 | TRUE |
| TYR | 393 | 25.51 | 9.76 | 0.68 | FALSE | 8.02 | 9.88 | 0.76 | TRUE |
| SER | 394 | 30.46 | 9.76 | 0.8 | FALSE | 16.26 | 13.53 | 0.51 | FALSE |
| VAL | 395 | 25.78 | 9.76 | 0.81 | FALSE | 11.58 | 13.53 | 0.93 | FALSE |
| LEU | 396 | 37.44 | 19.18 | 0.59 | TRUE | 13.95 | 12.5 | 0.7 | TRUE |
| TYR | 397 | 5.74 | 3.43 | 0.91 | TRUE | 53.46 | 9.95 | 0.76 | TRUE |
| ASN | 398 | 7.93 | 3.76 | 0.88 | FALSE | 34.82 | 6.63 | 0.73 | FALSE |
| SER | 399 | 8.79 | 3.76 | 0.91 | FALSE | 35.67 | 6.63 | 0.86 | FALSE |
| ALA | 400 | 6.56 | 3.76 | 0.91 | FALSE | 33.45 | 6.63 | 0.86 | FALSE |
| SER | 401 | 4.96 | 3.22 | 0.91 | FALSE | 33.85 | 6.63 | 0.88 | FALSE |
| PHE | 402 | 3.19 | 3.22 | 0.92 | FALSE | 7.12 | 4.92 | 0.87 | TRUE |

| | | | | | | | | | |
|---|---|---|---|---|---|---|---|---|---|
| SER | 403 | 32.56 | 7.15 | 0.82 | TRUE | 13.98 | 11.23 | 0.73 | TRUE |
| THR | 404 | 30.49 | 4.04 | 0.89 | TRUE | 20.18 | 16.25 | 0.64 | TRUE |
| PHE | 405 | 54.03 | 9.12 | 0.77 | TRUE | 51.4 | 10.85 | 0.74 | TRUE |
| LYS | 406 | 50.99 | 12.49 | 0.71 | TRUE | 17.14 | 6.41 | 0.84 | TRUE |
| CYS | 407 | 10.37 | 3.7 | 0.9 | FALSE | 54.79 | 9.61 | 0.76 | TRUE |
| TYR | 408 | 7.75 | 3.7 | 0.9 | FALSE | 50.73 | 12.27 | 0.71 | TRUE |
| GLY | 409 | 5.46 | 3.93 | 0.9 | TRUE | 59.74 | 6.99 | 0.81 | FALSE |
| VAL | 410 | 44.99 | 15.81 | 0.64 | TRUE | 56.6 | 6.99 | 0.82 | FALSE |
| SER | 411 | 8.49 | 3.46 | 0.91 | FALSE | 60.94 | 6.99 | 0.84 | FALSE |
| PRO | 412 | | | | | | | | |
| THR | 413 | 5.4 | 3.46 | 0.91 | FALSE | 23.5 | 3.92 | 0.9 | TRUE |
| LYS | 414 | 8.09 | 3.46 | 0.9 | FALSE | 49.66 | 12.05 | 0.71 | TRUE |
| LEU | 415 | 15.99 | 4.23 | 0.89 | TRUE | 52.52 | 10.22 | 0.75 | TRUE |
| ASN | 416 | 22.98 | 7.15 | 0.82 | TRUE | 28.97 | 8.4 | 0.7 | FALSE |
| ASP | 417 | 9.12 | 5.15 | 0.87 | TRUE | 29.78 | 8.4 | 0.87 | FALSE |
| LEU | 418 | 16.67 | 5.53 | 0.86 | TRUE | 23.05 | 8.4 | 0.83 | FALSE |
| CYS | 419 | 24.89 | 3.92 | 0.9 | FALSE | 29.71 | 8.4 | 0.78 | FALSE |
| PHE | 420 | 24.32 | 3.92 | 0.89 | FALSE | 20.26 | 9.97 | 0.76 | TRUE |
| THR | 421 | 39.16 | 8.11 | 0.78 | FALSE | 16.15 | 6 | 0.85 | TRUE |
| ASN | 422 | 43.16 | 8.11 | 0.81 | FALSE | 35.65 | 8.5 | 0.69 | FALSE |
| VAL | 423 | 37.05 | 8.11 | 0.75 | FALSE | 29.54 | 8.5 | 0.78 | FALSE |
| TYR | 424 | 36.88 | 8.11 | 0.85 | FALSE | 29.37 | 8.5 | 0.91 | FALSE |
| ALA | 425 | 49.15 | 15.47 | 0.65 | TRUE | 45.29 | 15.92 | 0.64 | TRUE |
| ASP | 426 | 38.91 | 6.82 | 0.77 | FALSE | 51.73 | 9.23 | 0.77 | TRUE |
| SER | 427 | 39.43 | 6.82 | 0.89 | FALSE | 29.86 | 4.5 | 0.88 | TRUE |
| PHE | 428 | 43.16 | 12.28 | 0.71 | TRUE | 15.1 | 5.99 | 0.85 | TRUE |
| VAL | 429 | 48.27 | 12.31 | 0.71 | TRUE | 21.1 | 7.63 | 0.81 | TRUE |
| ILE | 430 | 49.28 | 6.52 | 0.84 | FALSE | 48.28 | 11.88 | 0.72 | TRUE |
| ARG | 431 | 53.39 | 6.52 | 0.83 | FALSE | 50.37 | 11.09 | 0.73 | TRUE |
| GLY | 432 | 29.45 | 11.49 | 0.73 | TRUE | 25.67 | 9.75 | 0.88 | FALSE |
| ASP | 433 | 14.21 | 4.5 | 0.88 | TRUE | 26.14 | 9.75 | 0.66 | FALSE |

| | | | | | | | | | |
|---|---|---|---|---|---|---|---|---|---|
| GLU | 434 | 28.13 | 3.21 | 0.91 | TRUE | 9.57 | 6.4 | 0.84 | TRUE |
| VAL | 435 | 43.87 | 13.77 | 0.68 | TRUE | 44.26 | 13.88 | 0.68 | TRUE |
| ARG | 436 | 2.31 | 3.42 | 0.92 | FALSE | 41.42 | 8.78 | 0.72 | FALSE |
| GLN | 437 | 4.25 | 3.42 | 0.9 | FALSE | 43.36 | 8.78 | 0.85 | FALSE |
| ILE | 438 | 1.29 | 2.57 | 0.93 | TRUE | 45.07 | 13.34 | 0.69 | TRUE |
| ALA | 439 | 51.92 | 10.53 | 0.75 | TRUE | 16.72 | 5.07 | 0.87 | TRUE |
| PRO | 440 | | | | | | | | |
| GLY | 441 | 17.07 | 4.9 | 0.83 | FALSE | 10.2 | 4.64 | 0.83 | FALSE |
| GLN | 442 | 19.93 | 4.9 | 0.9 | FALSE | 13.05 | 4.64 | 0.9 | FALSE |
| THR | 443 | 19.36 | 4.9 | 0.9 | FALSE | 12.48 | 4.64 | 0.91 | FALSE |
| GLY | 444 | 19.59 | 4.9 | 0.92 | FALSE | 7 | 6.01 | 0.8 | FALSE |
| LYS | 445 | 19.36 | 4.9 | 0.82 | FALSE | 6.77 | 6.01 | 0.9 | FALSE |
| ILE | 446 | 30.62 | 18.85 | 0.59 | TRUE | 20.27 | 4.21 | 0.89 | TRUE |
| ALA | 447 | 26.95 | 21.89 | 0.54 | TRUE | 51.99 | 7.26 | 0.81 | FALSE |
| ASP | 448 | 50.56 | 11.43 | 0.73 | TRUE | 53.88 | 7.26 | 0.82 | FALSE |
| TYR | 449 | 51.96 | 11.27 | 0.73 | TRUE | 51.65 | 10.97 | 0.74 | TRUE |
| ASN | 450 | 51.31 | 11.63 | 0.72 | TRUE | 52.07 | 10.82 | 0.74 | TRUE |
| TYR | 451 | 51.34 | 10.79 | 0.74 | TRUE | 23.01 | 5.2 | 0.86 | TRUE |
| LYS | 452 | 44.11 | 7.56 | 0.83 | FALSE | 13.33 | 12.36 | 0.71 | TRUE |
| LEU | 453 | 41.42 | 7.56 | 0.86 | FALSE | 49.1 | 13.63 | 0.68 | TRUE |
| PRO | 454 | | | | | | | | |
| ASP | 455 | 43.26 | 7.56 | 0.74 | FALSE | 26.68 | 10.52 | 0.66 | FALSE |
| ASP | 456 | 5.56 | 3.83 | 0.9 | TRUE | 27.04 | 10.52 | 0.84 | FALSE |
| PHE | 457 | 11.53 | 1.78 | 0.95 | TRUE | 31.54 | 21.01 | 0.56 | TRUE |
| THR | 458 | 31.05 | 7.33 | 0.72 | FALSE | 33.35 | 12.2 | 0.68 | FALSE |
| GLY | 459 | 32.07 | 7.33 | 0.92 | FALSE | 34.38 | 12.2 | 0.74 | FALSE |
| CYS | 460 | 10.96 | 5.11 | 0.87 | TRUE | 34.15 | 15.02 | 0.61 | FALSE |
| VAL | 461 | 25.36 | 10.62 | 0.89 | FALSE | 28.78 | 15.02 | 0.71 | FALSE |
| ILE | 462 | 21.25 | 10.62 | 0.62 | FALSE | 26.02 | 8.01 | 0.71 | FALSE |
| ALA | 463 | 7.99 | 3.84 | 0.88 | FALSE | 29.67 | 8.01 | 0.85 | FALSE |
| TRP | 464 | 6.96 | 3.84 | 0.92 | FALSE | 28.65 | 8.01 | 0.84 | FALSE |

| | | | | | | | | | |
|---|---|---|---|---|---|---|---|---|---|
| ASN | 465 | 11.47 | 3.84 | 0.89 | FALSE | 31.59 | 6.92 | 0.85 | FALSE |
| SER | 466 | 13.24 | 3.84 | 0.91 | FALSE | 33.35 | 6.92 | 0.8 | FALSE |
| ASN | 467 | 33.42 | 9.79 | 0.79 | FALSE | 12.1 | 4.76 | 0.88 | TRUE |
| ASN | 468 | 33.54 | 9.79 | 0.78 | FALSE | 37.13 | 20.86 | 0.56 | TRUE |
| LEU | 469 | 27.43 | 9.79 | 0.71 | FALSE | 16.52 | 9.23 | 0.77 | TRUE |
| ASP | 470 | 16.4 | 3.97 | 0.89 | TRUE | 28.65 | 5.2 | 0.93 | FALSE |
| SER | 471 | 14.22 | 5.35 | 0.83 | FALSE | 30.37 | 5.2 | 0.75 | FALSE |
| LYS | 472 | 14.61 | 5.35 | 0.89 | FALSE | 30.75 | 5.2 | 0.93 | FALSE |
| VAL | 473 | 1.33 | 2.89 | 0.92 | FALSE | 25.96 | 5.2 | 0.86 | FALSE |
| GLY | 474 | 3.67 | 2.89 | 0.92 | FALSE | 40.3 | 26.04 | 0.48 | TRUE |
| GLY | 475 | 19.12 | 9.71 | 0.6 | FALSE | 25.86 | 4.27 | 0.89 | TRUE |
| ASN | 476 | 22.09 | 9.71 | 0.81 | FALSE | 19.37 | 11.12 | 0.73 | TRUE |
| TYR | 477 | 18.6 | 9.71 | 0.91 | FALSE | 4.87 | 3.12 | 0.92 | FALSE |
| ASN | 478 | 19.09 | 11.08 | 0.73 | TRUE | 7.67 | 3.12 | 0.92 | FALSE |
| TYR | 479 | 6.58 | 5.08 | 0.87 | TRUE | 4.87 | 3.12 | 0.91 | FALSE |
| LEU | 480 | 22.82 | 2.88 | 0.92 | TRUE | | | | |
| TYR | 481 | 37.27 | 22.49 | 0.53 | TRUE | 37.14 | 17.15 | 0.62 | TRUE |
| ARG | 482 | 40.72 | 13.37 | 0.75 | FALSE | 18.18 | 5.52 | 0.82 | FALSE |
| LEU | 483 | 37.92 | 13.37 | 0.63 | FALSE | 15.38 | 5.52 | 0.88 | FALSE |
| PHE | 484 | 19.46 | 5.08 | 0.87 | TRUE | 14.87 | 5.52 | 0.87 | FALSE |
| ARG | 485 | 27.55 | 3.11 | 0.92 | TRUE | 13.79 | 7.2 | 0.82 | TRUE |
| LYS | 486 | 4.99 | 3.35 | 0.91 | FALSE | 22.39 | 10.65 | 0.91 | FALSE |
| SER | 487 | 6.76 | 3.35 | 0.92 | FALSE | 24.16 | 10.65 | 0.73 | FALSE |
| ASN | 488 | 8.47 | 3.35 | 0.9 | FALSE | 25.87 | 10.65 | 0.62 | FALSE |
| LEU | 489 | 5.11 | 3.67 | 0.9 | TRUE | 34.32 | 12.02 | 0.66 | FALSE |
| LYS | 490 | 29.34 | 2.61 | 0.93 | TRUE | 34.38 | 12.02 | 0.78 | FALSE |
| PRO | 491 | | | | | | | | |
| PHE | 492 | 5.8 | 5.07 | 0.87 | TRUE | 20.49 | 4.09 | 0.9 | FALSE |
| GLU | 493 | 40.54 | 9.06 | 0.78 | TRUE | 22.96 | 4.09 | 0.9 | FALSE |
| ARG | 494 | 17.73 | 6.92 | 0.82 | TRUE | 22.85 | 4.09 | 0.88 | FALSE |
| ASP | 495 | 27.13 | 8.74 | 0.78 | TRUE | 28.08 | 4.31 | 0.89 | TRUE |

| | | | | | | | | | |
|---|---|---|---|---|---|---|---|---|---|
| ILE | 496 | 10.1 | 9.8 | 0.76 | TRUE | 14.86 | 4.17 | 0.89 | TRUE |
| SER | 497 | 17.18 | 5.56 | 0.86 | TRUE | 21.73 | 4.92 | 0.91 | FALSE |
| THR | 498 | 29.08 | 2.73 | 0.93 | TRUE | 22.25 | 4.92 | 0.84 | FALSE |
| GLU | 499 | 6.04 | 3.87 | 0.9 | TRUE | 21.76 | 11.63 | 0.72 | TRUE |
| ILE | 500 | 10.72 | 2.67 | 0.93 | TRUE | 43.31 | 18.92 | 0.59 | TRUE |
| TYR | 501 | 10.2 | 2.1 | 0.94 | TRUE | 5.19 | 5.59 | 0.65 | FALSE |
| GLN | 502 | 8.6 | 3.99 | 0.89 | TRUE | 8.67 | 5.59 | 0.89 | FALSE |
| ALA | 503 | 5.92 | 4.17 | 0.89 | TRUE | 9.18 | 5.59 | 0.9 | FALSE |
| GLY | 504 | 5.89 | 4.28 | 0.89 | FALSE | 7.87 | 5.59 | 0.91 | FALSE |
| SER | 505 | 9.14 | 4.28 | 0.87 | FALSE | 11.12 | 5.59 | 0.91 | FALSE |
| THR | 506 | 7.37 | 4.28 | 0.9 | FALSE | 9.35 | 5.59 | 0.91 | FALSE |
| PRO | 507 | | | | | | | | |
| CYS | 508 | 20.78 | 18.09 | 0.6 | FALSE | 8.4 | 4.05 | 0.89 | FALSE |
| ASN | 509 | 14.78 | 6.22 | 0.63 | FALSE | 12.17 | 4.05 | 0.89 | FALSE |
| GLY | 510 | 10.49 | 6.22 | 0.9 | FALSE | 7.89 | 4.05 | 0.9 | FALSE |
| VAL | 511 | 5.81 | 6.22 | 0.89 | FALSE | 19.11 | 8.56 | 0.77 | FALSE |
| GLU | 512 | 7.42 | 6.22 | 0.9 | FALSE | 20.72 | 8.56 | 0.59 | FALSE |
| GLY | 513 | 7.83 | 6.22 | 0.92 | FALSE | 21.13 | 8.56 | 0.91 | FALSE |
| PHE | 514 | 5.88 | 4.65 | 0.88 | TRUE | 21.73 | 8.56 | 0.92 | FALSE |
| ASN | 515 | 52.28 | 11.19 | 0.73 | TRUE | 6.72 | 4.35 | 0.89 | TRUE |
| CYS | 516 | 32.15 | 12.55 | 0.7 | TRUE | 25.2 | 13.19 | 0.64 | FALSE |
| TYR | 517 | 32.46 | 10.01 | 0.71 | FALSE | 21.43 | 13.19 | 0.75 | FALSE |
| PHE | 518 | 29.78 | 10.01 | 0.81 | FALSE | 22.56 | 8.54 | 0.79 | FALSE |
| PRO | 519 | | | | | | | | |
| LEU | 520 | 6.73 | 3.81 | 0.89 | FALSE | 37.53 | 6.29 | 0.76 | FALSE |
| GLN | 521 | 10.55 | 3.81 | 0.91 | FALSE | 41.35 | 6.29 | 0.86 | FALSE |
| SER | 522 | 43.83 | 12.29 | 0.74 | FALSE | 45.47 | 6.29 | 0.9 | FALSE |
| TYR | 523 | 40.74 | 12.29 | 0.68 | FALSE | 12.11 | 9.06 | 0.78 | TRUE |
| GLY | 524 | 18.2 | 13.43 | 0.69 | TRUE | 16 | 5.27 | 0.86 | TRUE |
| PHE | 525 | 4.34 | 3.37 | 0.91 | FALSE | 34.8 | 7.7 | 0.69 | FALSE |
| GLN | 526 | 5.42 | 3.37 | 0.91 | FALSE | 35.88 | 7.7 | 0.94 | FALSE |

| | | | | | | | | | |
|---|---|---|---|---|---|---|---|---|---|
| PRO | 527 | | | | | | | | |
| THR | 528 | 3.02 | 3.17 | 0.92 | TRUE | 17.34 | 2.3 | 0.94 | FALSE |
| ASN | 529 | 22.64 | 7.69 | 0.66 | FALSE | 36.56 | 10.68 | 0.7 | FALSE |
| GLY | 530 | 20.36 | 7.69 | 0.87 | FALSE | 34.28 | 10.68 | 0.69 | FALSE |
| VAL | 531 | 15.68 | 7.69 | 0.92 | FALSE | 29.59 | 10.68 | 0.85 | FALSE |
| GLY | 532 | 3.46 | 2.78 | 0.93 | TRUE | 26.11 | 10.44 | 0.63 | FALSE |
| TYR | 533 | 14.26 | 3.43 | 0.89 | FALSE | 26.51 | 10.44 | 0.82 | FALSE |
| GLN | 534 | 15.46 | 3.43 | 0.93 | FALSE | 27.71 | 10.44 | 0.81 | FALSE |
| PRO | 535 | | | | | | | | |
| TYR | 536 | 55.41 | 15.96 | 0.55 | FALSE | 21.39 | 13.35 | 0.54 | FALSE |
| ARG | 537 | 59.07 | 15.96 | 0.75 | FALSE | 25.04 | 13.35 | 0.89 | FALSE |
| VAL | 538 | 24.63 | 24.23 | 0.51 | TRUE | 6.59 | 6.25 | 0.84 | TRUE |
| VAL | 539 | 49.72 | 12.37 | 0.71 | TRUE | 35.94 | 8.47 | 0.79 | TRUE |
| VAL | 540 | 49.48 | 12.74 | 0.7 | TRUE | 47.64 | 12.77 | 0.7 | TRUE |
| LEU | 541 | 48.28 | 12.79 | 0.7 | TRUE | 46.2 | 12.98 | 0.7 | TRUE |
| SER | 542 | 31.52 | 3.16 | 0.92 | TRUE | 30.94 | 1.73 | 0.95 | TRUE |
| PHE | 543 | 52.43 | 9.95 | 0.76 | TRUE | 50.38 | 11.75 | 0.72 | TRUE |
| GLU | 544 | | | | | 11.27 | 4.01 | 0.91 | FALSE |
| LEU | 545 | 7.94 | 3.24 | 0.91 | TRUE | 7.4 | 4.01 | 0.88 | FALSE |
| LEU | 546 | 19.66 | 5.24 | 0.78 | FALSE | 4.12 | 4.86 | 0.87 | TRUE |
| HIS | 547 | 25.27 | 5.24 | 0.91 | FALSE | 16.68 | 5.06 | 0.87 | TRUE |
| ALA | 548 | 26.95 | 5.24 | 0.9 | FALSE | 30.4 | 38.33 | 0.34 | FALSE |
| PRO | 549 | | | | | | | | |
| ALA | 550 | 5.09 | 4.82 | 0.87 | TRUE | | | | |

### 3. Accessing data and running PFNet

#### Benchmark and experimental HX/MS datasets

The synthetic benchmark and experimental HX/MS datasets are provided in HXMS format, which includes isotopic mass envelope peaks or centroids for all labeling time points and peptides, defined by their start and end residues, along with key experimental parameters. The datasets are available at Zenodo (https://zenodo.org/records/17353731). The raw HX/MS data for T4 lysozyme are available from the PRIDE database (identifier PXD069693). The raw HX/MS data for ecDHFR and EEHEE_rd4_087 mini protein were previously reported (Lu *et al.*, *Nature Chemical Biology*, 2025, https://doi.org/10.1038/s41589-025-02049-1) and are available from the PRIDE database (identifier PXD057539).

The full experimental data in HXMS format are also provided as Supplemental Data for convenience.

#### Installing and running PFNet

PFNet accepts input files in HXMS format (Weber *et al.*, *bioRxiv*, 2025.10.14.682397), which can be generated using PFLink (https://huggingface.co/spaces/glasgow-lab/PFLink). PFLink supports data exported from major HX/MS analysis software packages, including BioPharma Finder (Fisher), HDExaminer (Trajan), DynamX (Waters), and HDX Workbench.

Code for dataset generation, model training, and inference, and access to pretrained models is available at: https://github.com/glasgowlab/PFNet. Installation instructions are provided in the *README.md* file within the same repository.

To facilitate broader accessibility, the trained PFNet model is also available as a cloud-based application on Hugging Face Spaces: https://huggingface.co/spaces/glasgow-lab/PFNet.
